# SRRM1 coordinates an alternative splicing program that promotes expression of oncogenic protein isoforms

**DOI:** 10.1101/2025.08.20.671097

**Authors:** Kamal Othman, Lauren Viola, Hira Fatima, Jessica Lapierre, Graham MacLeod, Craig D Simpson, Christopher Chu, Russell Beltran, Yangjing Zhang, Stephane Angers, Olivier Saulnier, C. Jane McGlade

## Abstract

The alternative splicing of the adapter protein NUMB is dysregulated in multiple cancer types, regulating its functional divergence towards either tumor suppression or oncogenesis in an isoform-dependant manner. Here we utilized a NUMB exon 9 (E9) splicing reporter in a genome-wide CRISPR screen to identify splicing regulators SRRM1 and SRSF11 that promote NUMB oncogenic splicing in colorectal, lung and breast cancer cell lines. We show that SRRM1 and SRSF11 share common protein interactors, RNA targets and effects on an oncogenic splicing program which favors the expression of pro-tumorigenic isoforms. In addition to NUMB E9, SRRM1 regulates cancer-associated splicing events in genes encoding signaling proteins, transcription factors and actin cytoskeleton regulators, many of which are also developmentally regulated, including CD44, MKNK2, ECT2, DIAPH1, KAT5, TCF7L2, FOXM1 and TBX3. Loss of SRRM1 in colorectal and lung cancer cells reduces cell proliferation, colony formation and invasion capabilities, as well as expression of tumour promoters Cyclin D1, Notum, and PRDX2. Our data indicate that SRRM1 regulation of alternative splicing represents a node to target multiple properties of malignant cells, with broad effects on cellular signaling, proliferation, EMT, apoptosis resistance and stemness.

## Introduction

Alternative splicing of pre-mRNA is a mechanism that contributes to proteomic diversity by generating mature mRNA transcripts that encode distinct protein products. The core of the splicing machinery is the spliceosome which consists of five snRNPs (small nuclear ribonucleoprotein particles; snRNP U1, U2, U4, U5 and U6) and hundreds of additional proteins that interact with cis elements at intron-exon boundaries to catalyze the splicing reaction ^1^. Additional cis elements act as splicing enhancers or splicing suppressors through the recruitment of auxiliary splicing regulators such as the SR and HNRNP protein families ^1–3^. Alternatively spliced exons frequently encode protein sequences within modular domains or intrinsically disordered regions that contain sites of post-translational modification or protein interaction motifs. Therefore, patterns of alternative splicing have important functional implications by re-wiring protein networks that impact cellular physiology ^3^ and contribute to both the transcriptomic and proteomic complexity that underlie cellular functioning and cell fate determination ^2^.

Mis-regulation of alternative splicing and the associated changes in protein isoform expression has been linked to cancer initiation, progression and drug resistance ^4^. Aberrant splicing can arise through multiple mechanisms, including mutations that disrupt cis-acting splice sites, or through mutations or misexpression of trans-acting splicing factors that alter spliceosome activity ^5^. Activation of oncogenic signaling pathways can also alter splicing factor activity to promote the reversal of developmentally regulated splicing patterns. This can result in the expression of tumorigenic protein isoforms and/or reduced expression of isoforms normally associated with cell differentiation and cell fate ^4^.

Alternative splicing of the NUMB gene is developmentally regulated and is frequently altered in cancer ^6–11^. In mammals, NUMB has an essential role during development and a conserved function as an inhibitor of Notch-dependent activation of gene transcription ^9,12,13^. Mammalian NUMB also functions to regulate activity of additional pathways including receptor tyrosine kinase, Hedgehog and TP53 signaling ^14–16^. NUMB is alternatively spliced to generate four protein isoforms with distinct patterns of expression in different cell types ^9,10^. The two largest isoforms (p71 and p72) arise due to inclusion of coding exon 9 (E9) that inserts 49 amino acids in the central region of the protein. The two smaller proteins (p65 and p66) result from E9 skipping. Expression of NUMB E9 is developmentally regulated with stem and progenitor cells primarily expressing E9 included isoforms and differentiated cells primarily expressing E9 skipped isoforms ^6,10,17–19^. A number of tissue-specific splicing factors have been shown to alter NUMB E9 splicing during development and cell differentiation including Rbfox3 ^18^, SRSF family members ^6,17,20^, RBM4/5/10 ^21^ and PTBP1 ^17,22^.

Dysregulation of NUMB alternative splicing is broadly associated with multiple cancer types, and is characterized by increased expression of E9 included isoforms that impact cellular processes associated with tumor progression ^7,9,11,15,23^. Despite being broadly implicated in multiple tumour types, the mechanisms that dysregulate NUMB E9 splicing in the context of cancer are less well understood. While mutations in the NUMB gene have not been identified, alterations in splicing factor activity have been implicated in lung cancer where loss of function mutations in RBM10, for example, were shown to increase E9 inclusion ^7^. Other evidence suggests that oncogenic signaling, particularly through RAS, alters NUMB splicing at least partly through PTBP1 expression ^17,22^.

Understanding the mechanistic details by which splicing alterations arise and contribute to cancer provides an opportunity to develop strategies to therapeutically modify pathological splicing events and/or the downstream consequences. Here we use a genome-wide CRISPR screen based on the NUMB E9 splicing event to identify SRRM1 and SRSF11 as common factors that regulate cancer-associated splicing of NUMB exon 9 in colorectal, lung and breast cancer cells. Furthermore, we show that in addition to NUMB, SRRM1 regulates a set of tumor-associated splicing events and promotes expression of oncogenic protein isoforms associated with cell proliferation, actin cytoskeleton dynamics, signaling and stemness.

## Results

### A genome wide CRIPSR screen identifies regulators of NUMB exon 9 splicing

A mini-gene reporter in which alternative splicing of NUMB E9 can be monitored by expression of EGFP or mCherry fluorophores was constructed by cloning EGFP and mCherry coding sequences in tandem downstream of NUMB genomic region including exons 8-10 and corresponding introns. A frameshift mutation was inserted into E9 to allow for translation of EGFP or mCherry fusions in alternate frames depending on the inclusion or skipping of E9 (Fig 1A). The frame shift insertion of two nucleotides (E9+2) did not alter NUMB reporter splicing relative to the non-mutagenized version of the reporter (Supp Fig 1A/B). Reporter responsiveness was validated based on expression of EGFP and mCherry fusions at both the mRNA and protein level in response to transient transfection with known NUMB splicing regulators, constitutively active MEKK (MEKK-EE) or splicing factor SRSF1 (Supp Fig 1C/D).

**Figure 1:**
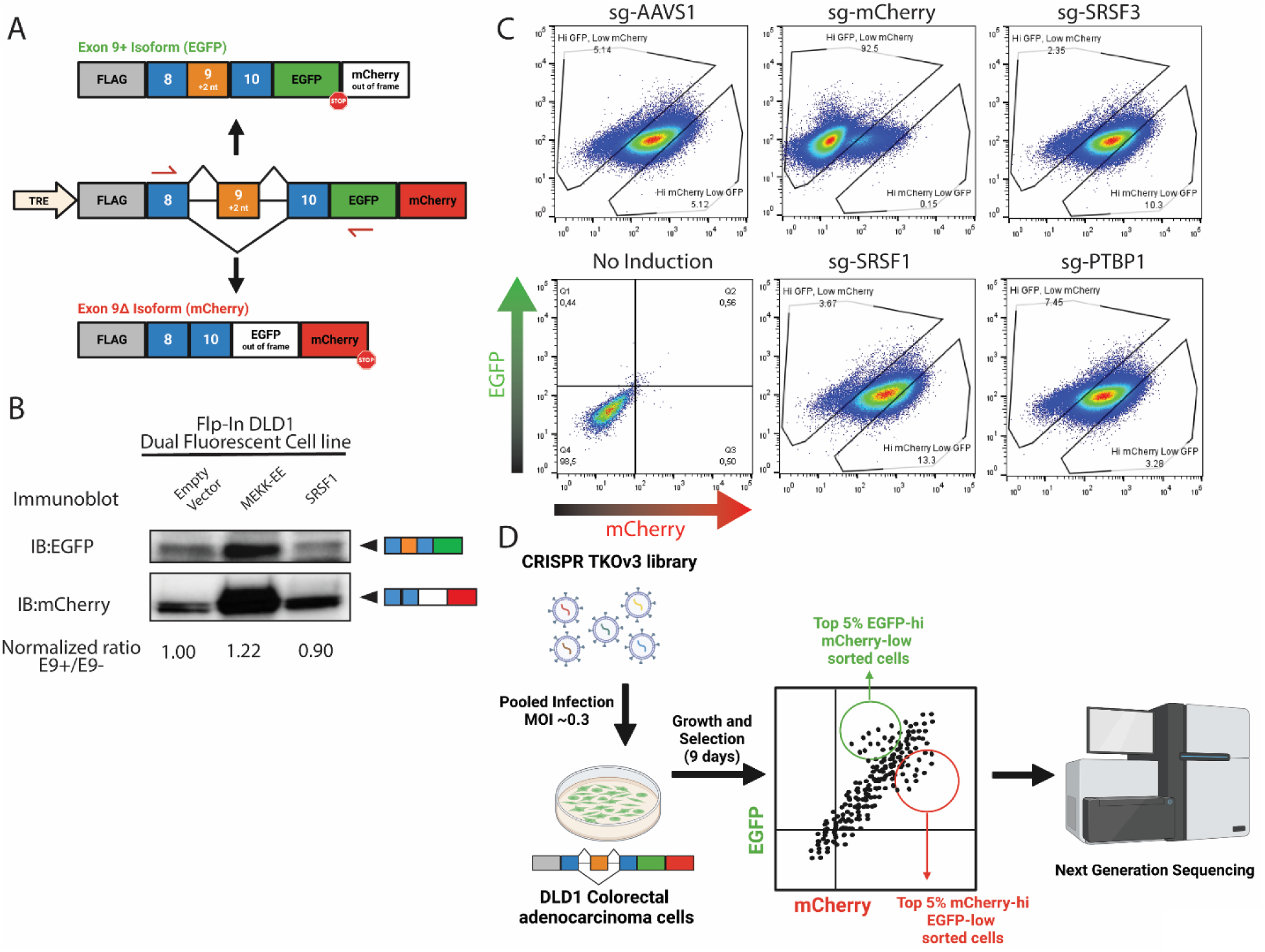
Dual fluorescent Numb splicing reporter based CRISPR knockout screen. **(A)** Schematic diagram of the dual fluorescent Numb E9 Splicing reporter. The reporter includes Numb exon 8, E9 (including a +2 bp frameshift) with E9 flanking introns (2934-nt upstream and 1987-nt downstream) and 88bps of exon 10. Mutations were introduced into exon such that the inclusion or skipping of E9 results in EGFP or mCherry expression respectively. **(B)** Immunoblots of lysates from E9 reporter expressing Flp-in DLD1 cells, transfected with MEKK-EE, SRSF1 or empty vector and probed with either EGFP a or mCherry antibodies. Numbers under blots denote ratio of relative band intensity quantifications utilizing ImageJ imaging software of the inclusion (EGFP)/exclusion (mCherry) product normalized to empty vector control. **(C)** FACS analysis of Flp-in DLD1 dual fluorescent reporter cell line following 24hr induction and transduced with lentiviral expression cassettes for Cas9 and sgRNAs targeting known RBP regulators of Numb E9. Gates were generated to capture the Top 5% EGFP high:mCherry low and EGFP low:mCherry high populations on the negative control sg-AAVS1 cells and were maintained for all the other targeted gene conditions. **(D)** Overview of genome-wide CRISPR knockout screen using the TKOv3 sgRNA library to identify novel pathways and factors regulating Numb E9 splicing. Dual fluorescent flp-in DLD1 Numb E9 reporter cell lines were transduced with a human CRISPR lentiviral library that has 70,948 guides (4 guides/gene) targeting 18,053 protein-coding genes at ∼0.3 MOI and subjected to puromycin selection for 9 days before PFA fixation and fluorescent activated cell sorting (FACs). Genomic DNA from the top 5%/20% EGFP high:mCherry low and mCherry-high:EGFP-low populations sorted populations was collected and sequenced via NGS.

An inducible DLD1 reporter cell line was generated (Fig 1B) and its suitability for a flow-based genome wide CRISPR screen was verified by transduction with lentiviral vectors expressing Cas9 and sgRNAs targeting known NUMB E9 splicing regulators SRSF1, SRSF3 or PTBP1 sgRNAs, as well as negative and positive controls targeting AAVS1 or mCherry respectively (Fig 1C). Reporter expression in transduced cells was monitored using fluorescence-activated cell sorting (FACS) and gated to capture the top 5% EGFP high/mCherry low and mCherry high/EGFP low populations. Knockout of PTBP1 or SRSF3 and SRSF1 resulted in a shift of the reporter population towards the EGFP high:mCherry low population or the EGFP low:mCherry high population respectively, confirming that the DLD1 reporter cell line was responsive to CRISPR-Cas9 deletion of splicing regulators (Fig 1C).

To identify factors that regulate NUMB E9 splicing, the DLD1 reporter cell line was transduced with the TKOv3 pooled sgRNA library targeting 18,053 protein-coding genes (4 guides per gene) and cultured for 9 days in puromycin selection. Cells within the top ∼5% or ∼20% of the EGFP high/mCherry low or EGFP low/mCherry high populations were captured by FACS and the sgRNA enrichment was analyzed via high throughput sequencing (Fig 1D). The data were collected from three independent biological replicates. The efficiency of sgRNA targeting and the effect of fitness on sgRNA enrichment scoring was tracked through the loss of sgRNA sequences in the unsorted cell populations at day 0 (T0, after selection) versus day 9 (T9, screen endpoint). Bayes factor (BF) scores ^24,25^ were determined using the BAGEL algorithm. Precision versus recall scores of core fitness DLD1 genes (Supp Fig 2A) ^25^ demonstrated efficient (AUC = 0.9704) sgRNA targeting considering the shorter screen duration (9 days). Bayes factor distribution of DLD1 non-essential versus fitness genes determined through BAGEL analysis (FDR < 0.05), yielded good separation indicating that CRISPR-Cas9 targeting was successful (Supp Fig 2B). In total, 515 unique hits were identified (false discovery rate FDR < 0.2, log fold enrichment LFC > 0.585) across both the 5% and 20% sorted populations (Supplementary Data 1).

**Figure 2:**
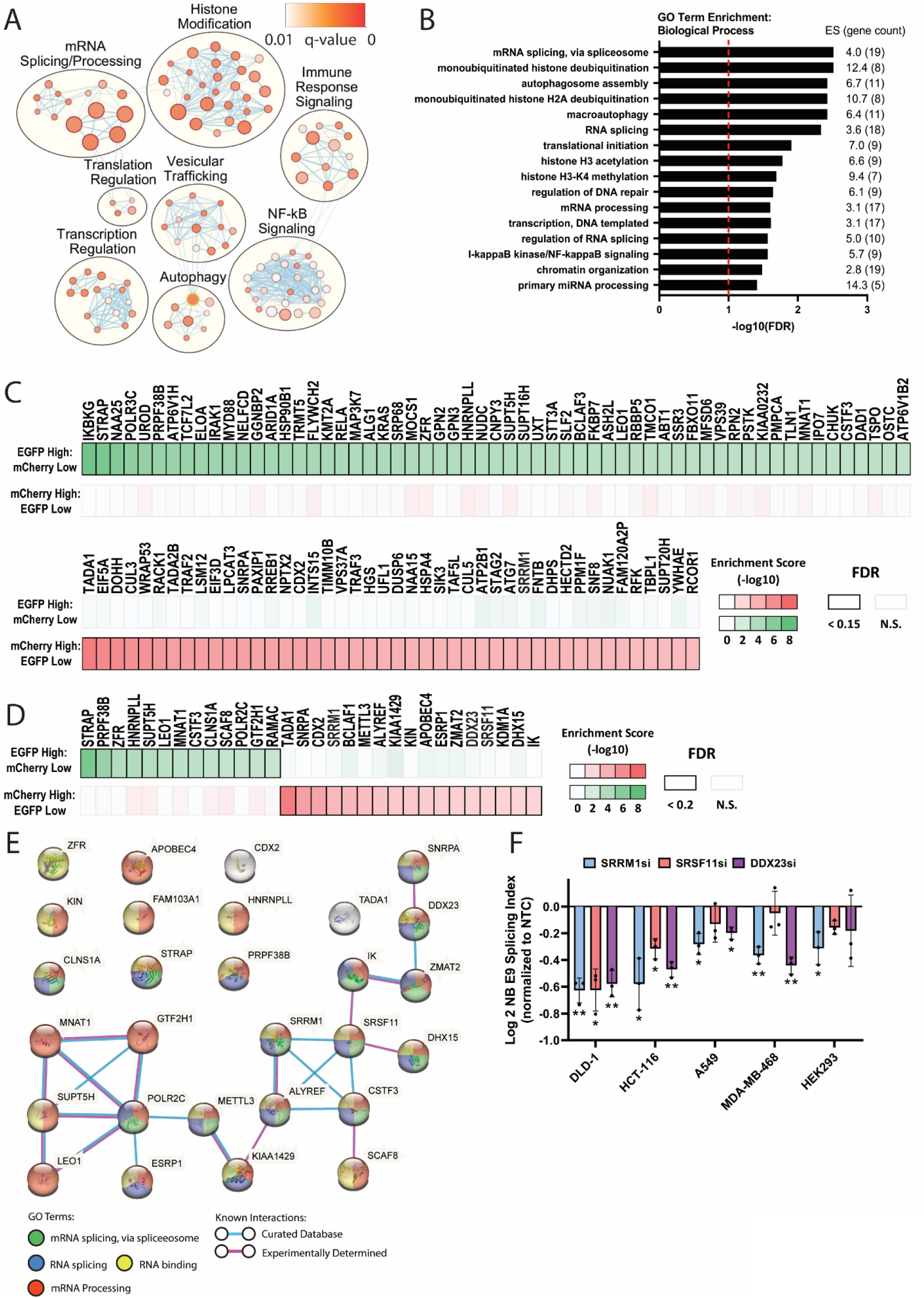
Identification of Numb exon 9 splicing reporter regulators. **(A)** Enrichment map of GO biological processes and reactome terms from the CRISPR screen hits (All unique hits from 5% and 20% sorted populations FDR <0.2, LFC >0.585). Only genes with an expression >1.0 TPM in DLD1 cells were considered. Node size represents the number of genes belonging to individual terms; color of node represents significance of the gene set (only q value < 0.01 gene sets are shown), edge thickness denotes shared interactors between terms. **(B)** Biological process GO term enrichment analysis for CRISPR screen hits. The same hit list as in **(A)** was uploaded to the online Database for Annotation, Visualization and Integrated Discovery, DAVID, and clustered using an EASE of 1.0 and medium stringency. The top enriched terms are arranged in order of significance (-log10(FDR) with Enrichment score (ES) values and total hit gene count for each term in between brackets denoted on the right. Only terms with an FDR score < 0.05 are shown. The red dotted line indicates an FDR of 0.1. **(C)** Relative effects of identified CRISPR screen hits detected by sgRNA enrichment in the 5% sorted cell populations. Genes identified in the 5% EGFP high/mCherry low sorted population (green) and the 5% mCherry high/EGFP low population (red) from DLD1 splicing reporter expressing cells with an FDR < 0.15, a log fold enrichment (LFC) > 0.585 and gene expression >1.0 TPM in DLD1 cells are shown. **(D)** List of CRISPR screen hits identified with a previously established link to splicing regulation with an FDR <0.2, LFC > 0.585 and TPM >1 in DLD1 cells from the 5% sorted populations. **(E)** STRING interaction network (https://string-db.org/) of genes shown in **(D)** identifying a potential regulatory splicing complex operating on Numb E9 alternative splicing **(F)** Examination of endogenous Numb E9 splicing index (log2 fold change expression of the Numb+E9 transcript relative to total Numb transcript expression, measured via RT-qPCR.) in DLD1, HCT-116, A549, MDA-MB-468 and HEK293 cells following transfection with siRNAs targeting SRRM1, SRSF11 or DDX23. (n = 3, mean Log2SI ± s.e.m.) All results depicted are relative to non-target control siRNA and significant differences with a p-values < 0.05 indicated with * and p-value < 0.01 with ** according to paired t-test statistical analysis.

The enriched biological process terms from Gene Ontology (GO) analysis of screen hits included those related to mRNA splicing/processing and histone modification (Fig 2A/B). Interestingly, other than the nucleoplasm which is the main site of splicing factor localization ^26^, cellular component GO enrichment analysis also identified the Spt-Ada-Gcn5 acetyltransferase (SAGA) complex which is an evolutionary conserved transcriptional coactivator that regulates gene expression through histone acetyltransferase and deubiquitylase activity (Supp Fig 3A). Among the identified genes, 47 are annotated in the Cancer Gene Census (CGC) according to the Catalogue of Somatic Mutations in Cancer (COSMIC, https://cancer.sanger.ac.uk/census) including several known oncogenic driver genes such as TP53, PIK3CA, KRAS, and ARID1A. Several splicing factors previously shown to regulate NUMB E9 AS were also identified including SRSF3 ^20^ and ESRP1 ^27^.

**Figure 3:**
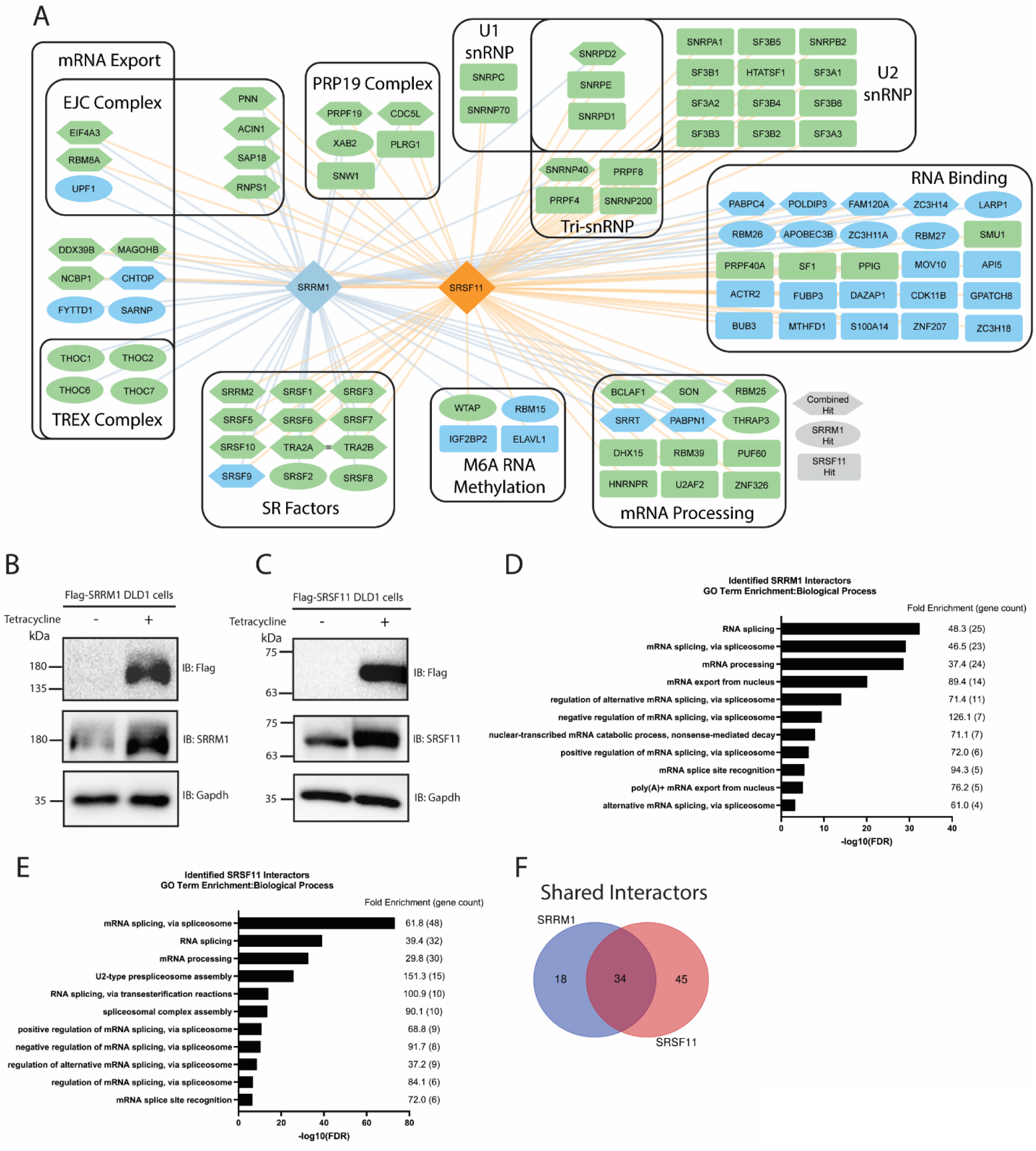
Investigating SRRM1/SRSF11 interactome using AP-MS in DLD1 colorectal adenocarcinoma cells. **(A)** Protein-Protein interactors of SRRM1 (blue edges) and SRSF11 (orange edges) identified via affinity-purification mass spectrometry in DLD1 cell lines. Interactors with a BFDR ≤ 0.05 are depicted. Unique prey are depicted in ovals for SRRM1 and rectangles for SRSF11 whilst shared prey between the two baits are depicted in hexagons. Green Nodes denote hits that possessed a GOBP term that belongs to RNA splicing and/or mRNA splicing, via spliceosome according to DAVID analysis (https://davidbioinformatics.nih.gov). **(B)** Immunoblots demonstrating expression of Flag tagged SRRM1 DLD1 cells ± tetracycline (10ug/ml) induction blotted with either anti-FLAG or anti-SRRM1 antibodies with corresponding anti-Gapdh blots used as a loading control. **(C)** Immunoblots demonstrating expression of Flag tagged SRSF11 DLD1 cells ± tetracycline (10ug/ml) blotted with anti-FLAG or anti-SRSF11 antibodies with corresponding anti-Gapdh blots used as a loading control. **(D)** Biological process GO term enrichment analysis for SRRM1 protein interactors analyzed through DAVID. The top enriched terms are arranged in order of significance (-log10 (FDR)) with fold enrichment values and total hit gene count for each term in between brackets denoted on the right. **(E)** Same as **(D)** for SRSF11 protein interactors. **(F)** Venn diagram depicting the number of shared interactors between SRRM1 and SRSF11 in DLD1 cells.

High confidence hits from the top 5% EGFP high or mCherry high sorted populations representing genes whose knockout had the largest effect on the splicing of the E9 reporter (Fig 2C) and hits with a previously established role associated with alternative splicing (Fig 2D) were prioritized for validation. High confidence hits identified included the splicing factors serine and arginine repetitive matrix 1 (SRRM1), serine and arginine rich splicing factor 11 (SRSF11) and DEAD-box helicase 23 (DDX23) whose involvement in regulating NUMB E9 AS has not previously been reported. String analysis of the splicing associated hits from the 5% sorted populations (Fig 2E) highlighted potential interactions of multiple factors identified from the screen, including connections between SRRM1, SRSF11, and DDX23.

### Splicing factors SRRM1, SRSF11 and DDX23 promote NUMB exon 9 inclusion

To validate potential NUMB E9 splicing regulators, we examined the effect of siRNA knockdown of a selected subset of 33 hits on endogenous NUMB E9 splicing. Selected targets whose knockdown resulted in a greater than 0.25 Log2PSI change compared to non-targeted siRNA controls were further validated in biological triplicate (Supp fig 3C). Amongst these, siRNA mediated knockdown of SRRM1, SRSF11 and DDX23 demonstrated strong and consistent effects on endogenous NUMB E9 splicing (Fig 2F). Targeting of gene expression was confirmed in siRNA treated cells via RT-qPCR and immunoblot (Supp fig 3 D/E). The effect of SRRM1, SRSF11 and DDX23 knockdown on endogenous NUMB E9 splicing was further examined in HEK293, MDA-MB-468 breast cancer cells and A549 lung cancer cells. SRRM1 knockdown with either siRNA or shRNA targeting resulted in greater E9 skipping across all cell lines tested (Fig 2F and Supp fig 3F/G), indicating a broad role for this factor in promoting inclusion of NUMB E9. Depletion of DDX23 and SRSF11 also resulted in E9 skipping across multiple cell lines tested but had inconsistent effects in HEK293 and MDA-MB-468 cells. These results identify SRRM1, DDX23 and SRSF11 as novel regulators of endogenous NUMB E9 splicing in multiple cellular contexts.

### SRRM1 and SRSF11 have common interactions with SR proteins and the splicing machinery

SRRM1 and SRSF11 are members of the SR family of splicing factors and their expression and activity have been implicated in cancer associated splicing events ^28–31^. To further elucidate the mechanisms underlying splicing regulation by SRRM1 and SRSF11 we used mass spectrometry to identify proteins that co-precipitated with Flag tagged SRRM1 or Flag tagged SRSF11 from DLD1 or HEK293T cell lysates (Supplementary data 2). The SRRM1 and SRSF11 interactomes were enriched for RNA binding proteins (RBPs), splicing factors and proteins involved in several spliceosome complexes including the core spliceosome snRNPs U1, U2 and tri-snRNP (U4/U5-U6) as well as the EJC, TREX and PRP19 complexes (Fig 3A, Supp Fig 4A). GO term enrichment analysis revealed a high enrichment of genes associated with RNA splicing via the spliceosome. (Fig 3D-E, Supp Fig 4D-E). Of note, SRRM1 was identified in the SRSF11 interaction network. While SRSF11 was not identified in the SRRM1 interactome, both proteins share the known SRRM1 binding partner, SRRM2, as well as multiple members of the SR family of splicing factors and components of the U1, U2 and Tri-snRNP ^32^. In agreement with prior studies demonstrating SRRM1 role in mRNA export by the Exon Junction Complex (EJC) ^33^ shared SRRM1 and SRSF11 interactors (Fig 3F) included EJC proteins EIF4A3, RBM8A and RNPS1, as well as factors also associated with mRNA export such as MAGOH, MAGOHB and DDX39B. In addition, multiple m6a mRNA methylation factors were also identified as SRRM1 interactors including WTAP, RBM15, ALKBH5 and ZC3H13 as well as VIRMA (also known as KIAA1429) (Fig 3A, Supp Fig 4A), which was also a hit in the splicing reporter screen (Fig 2D), while SRSF11 bound the m6a readers IGF2BP2 and ELAVL1 ^34^.

**Figure 4:**
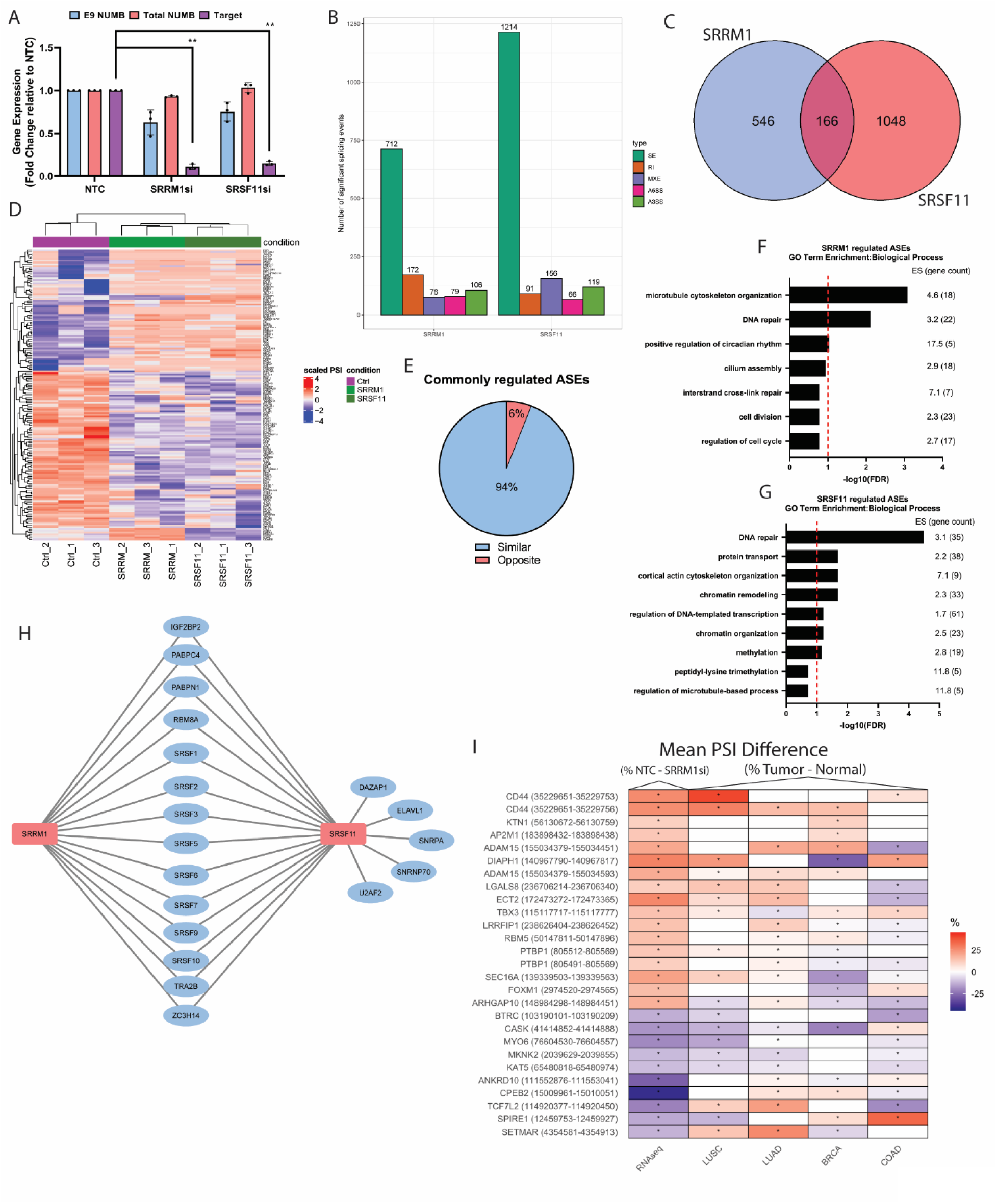
SRRM1 and SRSF11 knockdown results in altered downstream splicing. **(A)** Representative RT-qPCR of SRRM1 and SRSF11 silencing in DLD1 colorectal adenocarcinoma cell lines submitted for RNA-seq relative to non-target control. Error bars represent SEM (n =3 biological replicates for each condition, p-val < 0.001** according to paired student t-test). **(B)** Number of SRRM1 (left) or SRSF11 (right) dependent splicing events identified by rMATs following silencing treatment via siRNA. SE: alternatively spliced exon, IR: intron retention, MXE: mutually exclusive exons, A5SS: alternative 5’ splice sites, A3SS: alternative 3’ splice sites. **(C)** Venn diagram depicting overlap in SRRM1 and SRSF11 regulated alternatively spliced exon (ASE) events. **(D)** Heatmap of normalized PSI values of ASE events regulated by both SRRM1 and SRSF11 in siSRRM1, siSRSF11 and siCTRL samples. **(E)** The proportion of common target ASEs following SRRM1 or SRSF11 silencing shown in (C) (n=166) that are “similar” (blue) or “opposite” (orange) depending on whether delta PSI values vary in the same or opposite direction relative to control PSI values. **(F)** Biological process GO term enrichment analysis for all genes that demonstrated dysregulated alternatively spliced exons (ASEs) regulated by SRRM1 analyzed through DAVID. The top enriched terms are arranged in order of significance (-log10(FDR) with Enrichment score (ES) values and total hit gene count for each term in between brackets denoted on the right. The red dotted line indicates an FDR of 0.1. **(G)** Same as (**F**) for the list of genes that demonstrated dysregulated ASEs regulated by SRSF11. **(H)** Depicts proteins that were identified as protein-protein interactors of SRRM1 and/or SRSF11 through APMS with binding motifs enriched in the differentially spliced events downstream of SRRM1 and SRSF11 knockdown. **(I)** Heatmap depicting splicing events that demonstrated high correlation in mean PSI difference (%NTC-SRRM1si) in the DLD1 RNAseq splicing analysis compared to mean PSI difference (%Tumor – Normal) of LUSC, LUAD, BRCA and COAD TCGA patient samples. Only values that demonstrated a significant change in tumor vs normal patient samples are depicted.

### Identification of SRRM1 and SRSF11 regulated splicing events

To identify additional splicing events regulated by SRRM1 and SRSF11, we conducted bulk RNA-sequencing (RNA-seq) in DLD1 cells line following siRNA silencing of SRRM1 or SRSF11 (Fig 4A). Analysis of RNA-seq data demonstrated expected changes in NUMB E9 splicing (Supp Fig 5A/B). Principal component analysis of 3 biological replicates collected for each condition demonstrated high similarity (Supp Fig 5C). Exonic splicing events with a ΔPSI (Percent Spliced In) > 10% and a FDR < 0.05 compared to control samples were considered alternatively spliced exon (ASE/SE) events of interest (Fig 4B, Supplementary data 3). In total, SRRM1 and SRSF11 silencing resulted in the identification of 712 and 1214 differential ASE events respectively, of which 166 events were common to both SRRM1 and SRSF11 (Fig 4C). Directionality of changes in response to SRRM1 or SRSF11 silencing of common ASEs was ∼94% concordant (156/166) with only 10 ASEs demonstrating opposite PSI shifts between the two splicing factors (Fig 4D/E, Supp Fig 5D/E/F). SRRM1 silencing resulted in both exon inclusion (406 events) and exon skipping (306 events), while SRSF11 silencing resulted in exon inclusion in 90 events and exon skipping in 60 events (Supp Fig 5D/E/F).

**Figure 5:**
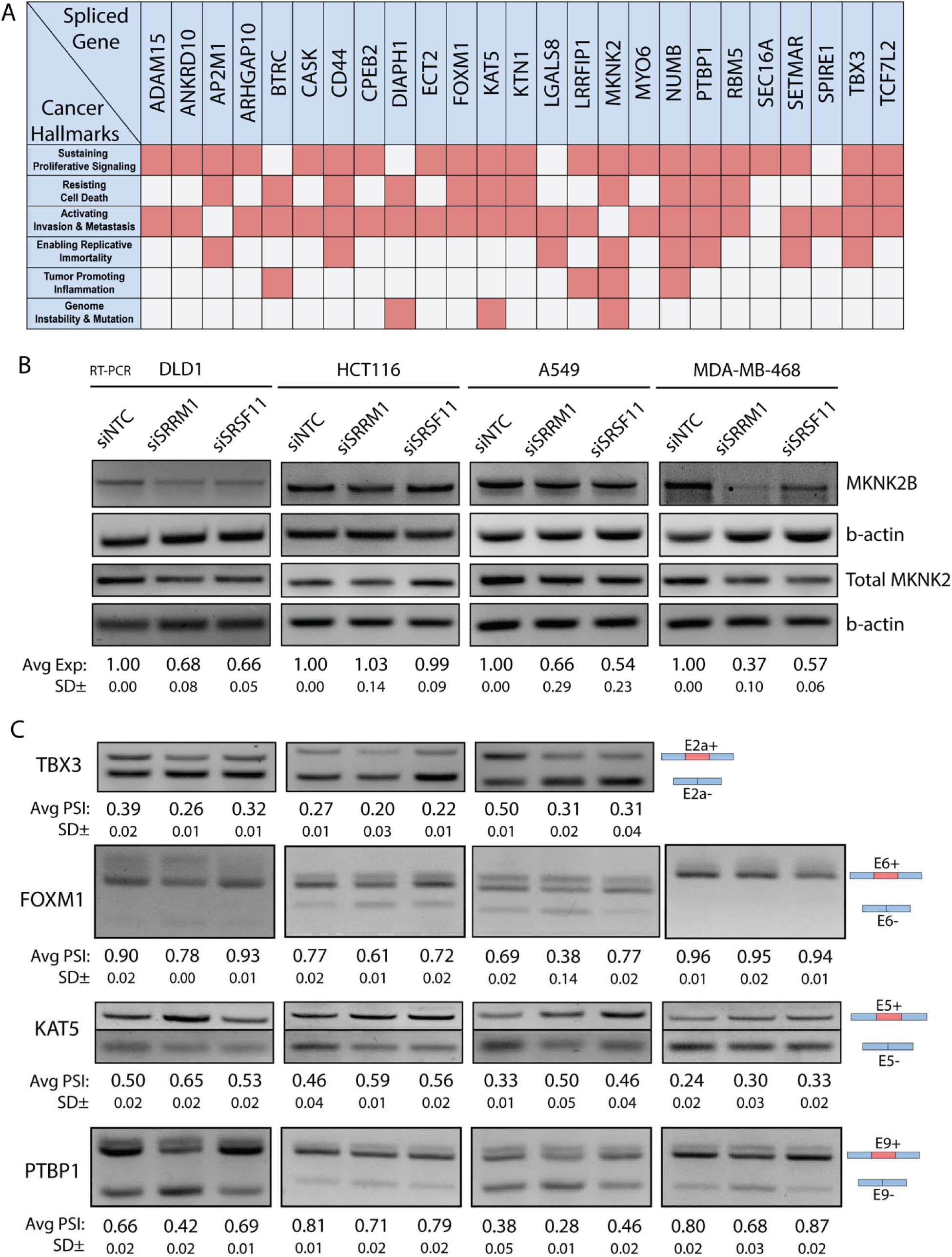
SRRM1 and SRSF11 promote expression of oncogenic protein isoforms in multiple cancer types. **(A)** Cancer hallmarks associated with genes that demonstrate dysregulated splicing upon SRRM1 knockdown identified through RNA-seq rMATs analysis **(B)** Representative semi-quantitative RT-PCR blots demonstrating effects of SRRM1 or SRSF11 knockdown via siRNA treatment on Total MKNK2 and MKNK2B isoform expression relative to β-actin compared with non-target control (NTC) treatment in DLD1, HCT116, A549 and MDA-MB-468 cancer cell lines. Average MKNK2B expression normalized to total MKNK2 expression ± SD of 3 biological replicates in each cell line is denoted below. **(C)** Representative semi-quantitative RT-PCR blots demonstrating effects of SRRM1 or SRSF11 knockdown via siRNA treatment on shifts in TBX3 exon 2a, FOXM1 exon 6, KAT5 exon 5 and PTBP1 exon 9 isoform expression compared with non-target control (NTC) treatment in DLD1, HCT116, A549 and MDA-MB-468 cancer cell lines. Average percent spliced in (PSI) ± SD of 3 biological replicates in each cell line is denoted below.

Comparison of differentially spliced genes (DSGs) with ASE events regulated by SRRM1 or SRSF11 revealed additional common targets shared between the two factors independent of exon location. These results suggest that SRRM1 and SRSF11 may regulate the splicing of different exons within the same gene (Supp Fig 5G). GO term analysis of the biological process enriched terms associated with the DSGs included DNA repair (Fig 4F/G) in both SRRM1 and SRSF11 knockdown cells, as well as microtubule cytoskeleton organization (SRRM1 silencing) and actin cytoskeleton organization (SRSF11 silencing) (Fig 4F/G).

Analysis of altered gene expression in response to SRRM1 and SRSF11 silencing revealed only 78 differentially expressed genes (DEGs) in SRRM1 depleted cells and 206 DEGs in SRSF11 depleted cells (fold change > 2, FDR < 0.05)(Supplementary data 3). Among the DEGs identified, 7 were shared between SRRM1 and SRSF11, 5 of which were altered similarly in both conditions (Supp Fig 5H). Comparison of the DSGs and DEGs revealed minimal overlap with only 3 genes common between the DSGs and DEGs with SRRM1 and SRSF11 silencing (Supp Fig 5I/J). This suggests that silencing of SRRM1 and SRSF11 primarily affects alternative splicing events, and secondarily alters the expression of a specific subset of genes.

Enriched RBP motifs near the SRRM1 and SRSF11 regulated ASEs were identified using rMAPS2 (Supp Fig 6). The majority of the top 20 enriched motifs identified were common to both SRRM1 and SRSF11(Supp Table 1and 2). While SRRM1 does not have well defined target RNA binding motif, among the most highly enriched motifs identified in both the SRRM1 and SRSF11-regulated ASEs were the SRSF10 binding motifs A[AG]AG[AG][AG][AG] and AGAGA[AG][AG]. Of note these motifs are highly similar to the less well characterized SRSF11 binding motif AG[AG][AG][AG] providing further evidence that SRSF11 likely regulates several of the ASEs also regulated by SRRM1 (Fig 4C/D/H). RBPs corresponding to 19 of the most enriched motifs were also identified as SRRM1 and SRSF11 interactors (Fig 4H). These factors include members of the SR family of splicing factors (SRSF 1-3, 5-7, 9-10) as well as a core component of the EJC RBM8A, the mRNA processing factors PABPC4, PABPN1 and ZC3H1, the m6a reader IGF2BP2 and the oncogenic splicing factor TRA2B (Fig 4H). ASEs were also enriched for the epithelial splicing regulatory protein 1 (ESRP1) motif (Supp Fig 6A/C) in agreement with ESRP1 known role in NUMB E9 splicing ^27^.

**Figure 6.**
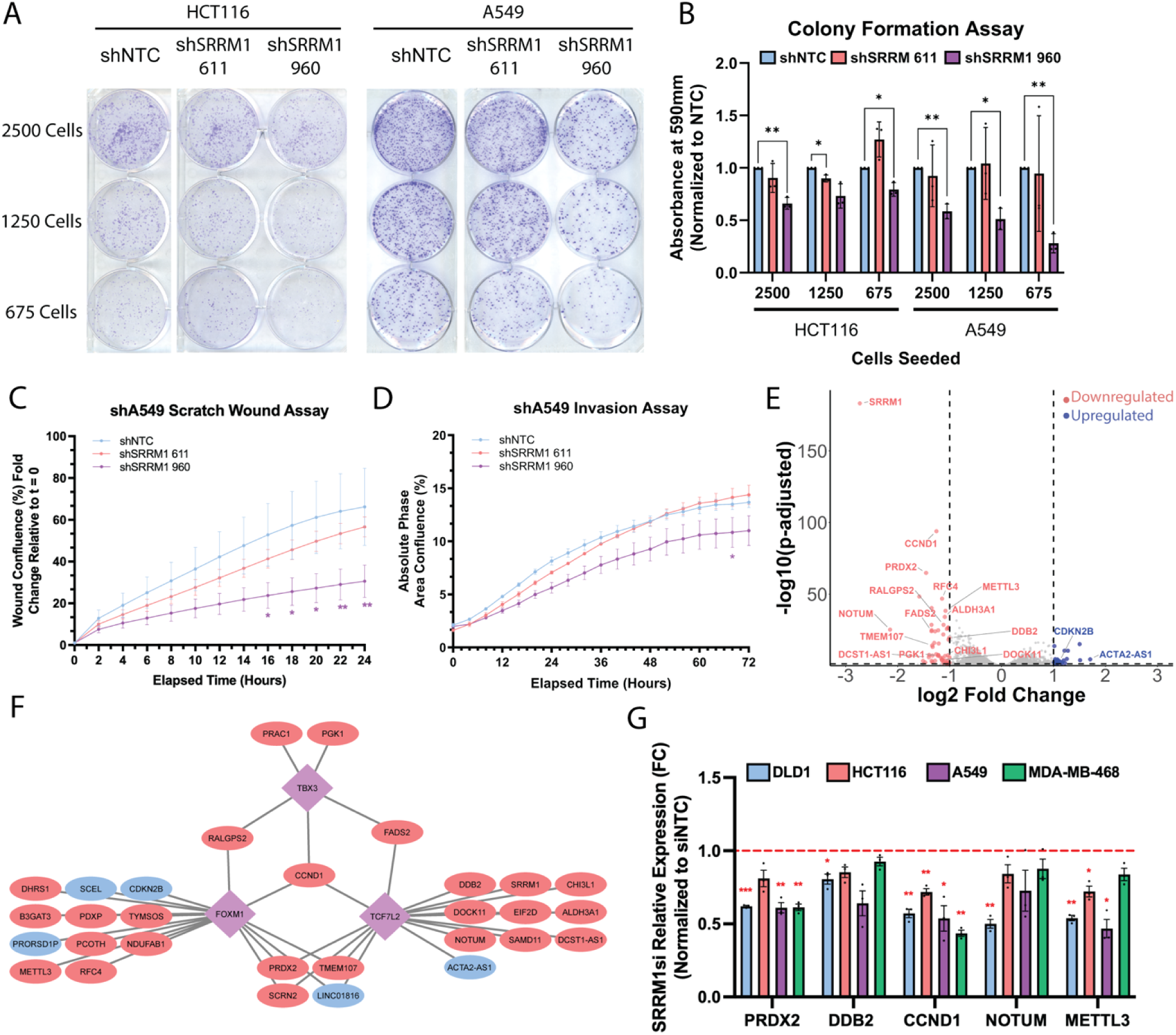
SRRM1 knockdown impairs cell growth and colony formation and alters gene expression. **(A)** Representative images from colony formation assays conducted on tetracycline-inducible HCT116 and A549 SRRM1 short hairpin cell lines (shSRRM1 611/960) relative to a non-target control line (NTC) seeded at either 2500, 1250 and 675 cells per well and cultured for 14 days (HCT116) or 10 days (A549) under tetracycline induction before crystal violet staining and imaging. **(B)** Quantification of colony formation assays **(C)** across 3 biological replicates normalized to NTC control. Significant differences are denoted with a p-values < 0.05 = *, p-values < 0.01 = ** according to a paired t-test. **(C)** Quantification of scratch wound migration assays conducted on A549 short hairpin cell lines measuring wound confluence using an Incucyte live cell imaging system. The wound confluence at each time point was normalized to the initial wound confluence. Cells were allowed to form a monolayer before scratching a wound and imaging migration across 24 hours. Significance was determine through a two-way ANOVA analysis between shNTC and shSRRM1 lines. **(D)** Same as **(C)** but cells were embedded in 8mg/ml Matrigel for an Invasion assay and imaged across 72 hours. **(E)** Volcano plot demonstrating -log10(FDR adjusted p-value) vs Log2fold change of gene expression in SRRM1 knockdown cells identified by RNA-seq Deseq2 analysis. Only genes that exhibited p-adjusted > 0.05 and Log2Fold change > 1 or < -1 were considered hits. Labeled downregulated oncogenic genes and upregulated tumor suppressive genes and potential targets of alternative spliced transcription factors (FOXM1, TBX3, TCF7L2) are shown. **(F)** Predicted targets of FOXM1, TBX3 and TCF7L2 transcription factors predicted through ChEA3 (https://maayanlab.cloud/chea3/), which exhibited significant expression changes in response to SRRM1 knockdown. Red nodes denote reduced expression, and blue nodes denote increased expression in response to SRRM1si treatment. **(G)** Real-time quantitative PCR measuring PRDX2, DDB2, CCND1, NOTUM and METTL3 gene expression in DLD1, HCT116, A549 and MDA-MB-468 cells treated with SRRM1 siRNA relative to non-target control treatment across 3 biological replicates. Relative fold change expression values were normalized to NTC control (red dotted line FC =1.0). Significant differences with a p-values < 0.05 = *, p-values < 0.01 = **, p-values < 0.001 = ***.

**Table 1:**
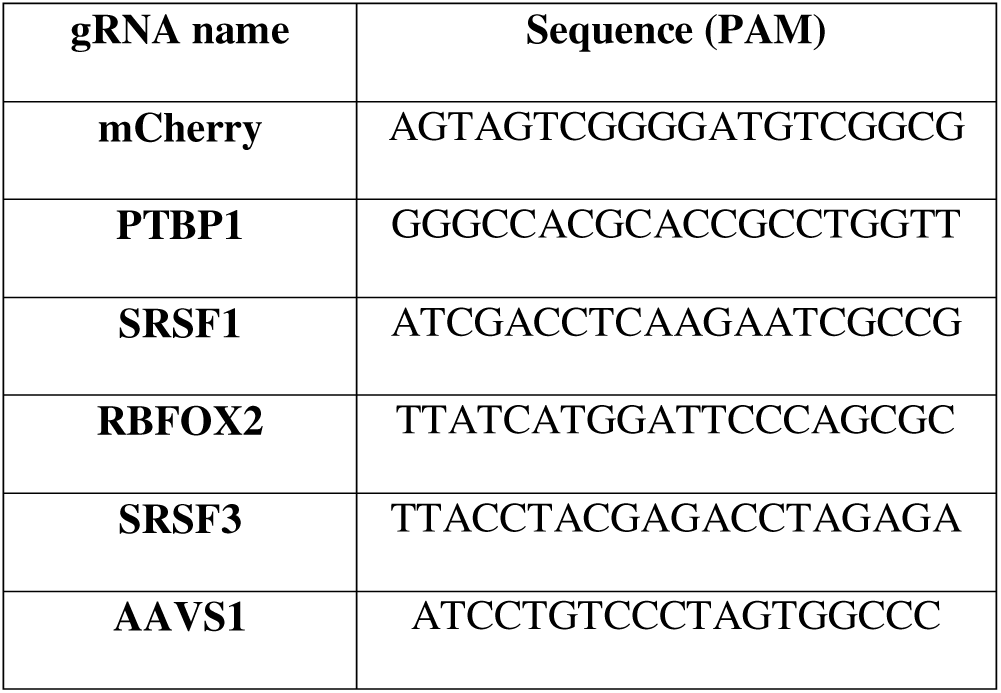
gRNA sequences used in splicing factor KO FACs test screen. Related to. **Figure 1C**.

### SRRM1 and SRSF11 promote a set of cancer-associated splicing events

In order to identify SRRM1 and/or SRSF11 regulated splicing events that may contribute to oncogenesis, we analyzed the list of differentially spliced genes (DSGs) for significant PSI changes in The Cancer Genome Atlas (TCGA) patient samples ^35^. Of the 593 SRRM1-regulated DSGs identified, 388 exhibited significant splicing changes in COAD, 479 in BRCA, 413 in LUAD and 457 in LUSC patient samples (Supp Fig 7A). Similarly, of the 949 SRSF11 regulated DSGs, 642 exhibited significant PSI changes in COAD, 788 in BRCA, 633 in LUAD and 753 in LUSC patient samples (Supp Fig 8A). Next, we examined how specific shifts in exon PSI varied with SRRM1 silencing (Supp Fig 7B-E) or SRSF11 silencing (Supp Fig 8B-E) in comparison to exon-matched PSI shifts in COAD, LUAD and LUSC tumor and normal patient samples. We found that 40∼41% of SRRM1 and 41∼47% of SRSF11 exon-matched ASEs that exhibited reduced PSI with factor knockdown, had a higher PSI in tumors versus normal TCGA patient samples or vice versa (Supp Fig 7B-E, 8B-E).

Of the SRRM1 regulated splicing events that are also altered in COAD, BRCA, LUAD or LUSC patient samples, we identified a set of 22 events that are annotated in VastDB and predicted to impact protein isoform expression (Fig 4I). The set of splicing events impact genes with known functions related to the hallmarks of cancer including proliferation, invasion and metastasis (Fig 5A). For example, the splicing of CD44 which has a well characterized role in tumour invasion and metastasis ^36,37^ has also previously been identified as an SRRM1 target ^38^. We next examined a subset of these cancer associated splicing events in colon, lung and breast cancer cell lines with SRRM1 or SRSF11 siRNA knockdown (Supp Fig 9A/B). MKNK2 is a serine threonine kinase that undergoes alternative splicing of exon 14a to form the functionally distinct isoforms MKNK2A that acts as a tumor-suppressor by inducing p38-mediated cell death and MKNK2B which activates eIF4E without triggering the p38 MAPK stress response ^39,40^. Analysis of TCGA RNA-seq data demonstrates that MKNK2 exon 14a PSI is significantly lower in COAD, LUAD and LUSC tumours (Supp Fig 9A). In line with these findings, we demonstrate that knockdown of SRRM1 or SRSF11 in DLD1, A549 and MDA-MB-468 cancer cell lines resulted in reduced MKNK2B isoform expression (Fig 5B and Supp Fig 10A).

SRRM1 knockdown also induced skipping of cassette exons in the T-box transcription factor (TBX3) as well as in the Forkhead box protein M1 (FOXM1) transcription factor. TBX3 is a transcriptional regulator linked to stem cell self-renewal and maintenance, and is a downstream effector of the Wnt signaling pathway ^41^. Both SRRM1 and SRSF11 knockdown reduced TBX3 exon 2a inclusion in DLD1, HCT116, and A549 cell lines (Fig 5C, Supp Fig 10B). TBX3 exon 2a is located within the T-box DNA binding domain, and skipping of this exon has been shown to be important for differentiation of embryonic stem cells ^42^,while inclusion in cancer is associated with senescence inhibition ^43^. In keeping with a tumour promoting role, exon 2a inclusion is significantly increased in BRCA, COAD, LUSC as well as BLCA and READ tumour types (Supp Fig 9A). Similarly, the Forkhead (FH) transcription factor, FOXM1 has been implicated in oncogenesis due to its elevated expression across several cancers as well as its ability to promote proliferation, metastasis, stemness and drug resistance ^44^. SRRM1 knockdown reduced the inclusion of FOXM1 exon 6 (also known as exon Va) in DLD1, HCT116 and A549 cancer cell lines (Fig 5C, Supp Fig 10C). FOXM1 exon 6 is located immediately adjacent to the FH DNA binding domain and contains a putative ERK1/2 phosphorylation site that renders the FOXM1c isoform MEK responsive ^45^. Analysis of TCGA data showed that exon 6 inclusion was significant higher in COAD (Supp Fig 9A) and prior studies have shown increased inclusion of exon 6 in pancreatic and ovarian tumours ^46,47^.

SRRM1-regulated exon matched splicing events that demonstrated significant PSI shifts across TCGA samples also included the histone acetyltransferase KAT5 (also known as TIP60) and the splicing factor PTBP1. KAT5 can act as a tumor suppressor by acetylating and activating pro-apoptotic genes such as p53 ^48,49^. The KAT5 exon 5 skipped isoform lacks the N-terminal domain required for chromatin binding and acts as a dominant negative inhibitor of the exon 5 included tumour suppressive isoform ^50^. In patient samples, KAT5 exon 5 PSI values were significantly lower in COAD, LUAD, LUSC and several other tumour types (Supp Fig 9A). Consistent with these results, knockdown of SRRM1 resulted in increased KAT5 exon 5 inclusion in DLD1, HCT116 and A549 cell lines. Similarly, SRSF11 knockdown had the same effect of increased exon 5 inclusion in HCT116 and A549 cell lines (Fig 5C, Supp Fig 10D).

SRRM1 knockdown also downregulated exon 9 inclusion of the hnRNP family member PTBP1 in all cell lines tested (Fig 5C, Supp Fig 10E). Increased inclusion of exon 9 was observed in both LUAD and THCA patient samples compared to normal control (Supp Fig 9A). Whilst the role of PTBP1 exon 9 as it relates to oncogenic activity has not been elucidated, exon 9 is important for regulation of PTBP1 splicing activity during neural development ^51^. Of note, PTBP1 has been shown to promote oncogenic splicing of both NUMB and PKM2 in breast cancer ^17,52^, as well as promote the stemness in gastric cancer ^53^.

We also identified previously uncharacterized splicing events in the Rho GEF epithelial cell transforming sequence 2 (ECT2) and Rho effector DIAPH1 that are regulated by SRRM1^54–56^. Knockdown of SRRM1 resulted in reduced ECT2 exon 4 inclusion in DLD1, HCT116, A549 and MDA-MB-468 cancer cell lines (Supp Fig 10F, Supp Fig 11F). Similarly, knockdown of SRSF11 also significantly reduced ECT2 exon 4 inclusion in A549 and MDA-MB-468 cells (Supp Fig 10F, Supp Fig 11F). Though its functional role has not been characterized, ECT2 exon 4 PSI is significantly higher in both LUAD and LUSC cancer samples (Supp Fig 9A). SRRM1 knockdown also resulted in reduced DIAPH1 exon 2 inclusion across DLD1, HCT116, A549 and MDA-MB-468 cancer cell lines whilst SRSF11 knockdown resulted in reduced DIAPH1 exon 2 inclusion only in the DLD1 and MDA-MB-468 cancer cell lines (Supp Fig 10G, Supp Fig 11G). A significant increase in DIAPH1 exon 2 PSI is observed in COAD, LUSC, and HNSC tumours (Supp Fig. 9). Taken together, these results demonstrate that SRRM1 and SRSF11 promote the expression of cancer-associated isoforms of multiple genes linked to multiple hallmarks of cancer.

### SRRM1 knockdown impairs cancer cell proliferation, colony formation and gene expression

To characterize the phenotypic effects of SRRM1 knockdown, we generated inducible short hairpin RNA (shRNA) HCT116 and A549 cell lines (Supp Fig 12A/B). Knockdown of SRRM1 resulted in significantly reduced colony formation (Fig 6A) and proliferation (Supp Fig 12D/E). SRRM1 knockdown also resulted in reduced cell migration as well as invasive capacity of A549 cell lines (Fig 6C/D). To assess how specific SRRM1 splicing events may contribute to the observed growth suppression in SRRM1 knockdown conditions, we considered the predicted downstream consequences of specific protein isoform expression. Silencing of SRRM1 results in altered splicing of MKNK2 and reduced expression of the MNK2B isoform which in turn activates phosphorylation of p38 and increases COX2 expression as part of the stress response pathway. Indeed, we observed enhanced phosphorylation of p38 and increased expression of p38 target gene COX2 in response to SRRM1 knockdown in DLD1, A549 and MDA-MB-468 cells compared to controls (Supp Fig 12F/G).

SRRM1 depletion also caused the altered expression of a small set of genes that are significantly enriched for E2F targets and G2M checkpoint associated genes (Supp Fig 5H,K; Supplementary data 3). In particular, SRRM1 knockdown in DLD1cells significantly reduced expression of genes with known functions in cell cycle progression (CCND1, RalGPS2), tumour progression and drug resistance (ALDHA1, PRDX2, PGK1), epithelial-mesenchymal transition (TMEM107, Lnc01816, DCST1-AS1), fatty acid metabolism (FADS2) and Wnt signaling (Notum, PRDX2, DDB2, CCND1, CDKN2B) (Fig 6E). Analysis of transcription factor target enrichment using ChEA3 ^57^ identified the SRRM1 splicing targets TBX3, FOXM1 and TCF7L2 amongst the most highly enriched for this differentially expressed gene set (Fig 6F). Analysis of selected target genes by RT-qPCR in DLD1, HCT116, A549 and MDA-MB-468 cells demonstrated reduced expression of PRDX2, DDB2, CCND1, NOTUM as well as METTL3 in response to knockdown of SRRM1 (Fig 6G). These data suggest that in addition to the oncogenic effects of directly regulating isoform expression of proteins such as NUMB, MKNK2 and CD44, SRRM1 can also promote a tumorigenic gene expression profile through altered splicing of transcription factors.

## Discussion

Detailed studies of the mechanisms and targets of dysregulated splicing can advance our understanding of tumourigenesis^4^. Here we used a CRISPR-Cas9 screen, based on a splicing event frequently dysregulated in cancer, to identify the SR protein family members, SRRM1 and SRSF11 as regulators of NUMB E9 splicing. We assessed the extent of functional overlap between SRRM1 and SRSF11 by comparing their shared interactomes and global effects on alternative splicing, as well as identify a set of coordinately regulated splicing events. SRRM1 functions as a splicing co-activator through association with other SR family members known to bind exonic splicing enhancers (ESEs). Indeed, SRRM1 and SRSF11 commonly regulated exons were enriched with binding motifs for an overlapping set of SR proteins including those that have been linked to cancer-associated splicing events including SRSF1, SRSF2, SRSF3, SRSF10 and TRA2B ^4^. This set of proteins were also present in the SRRM1 and SRSF11 interactomes, along with SRRM2 and the U1 and U2 snRNP spliceosome components that together form a splicing co-activator complex ^58^.

The roles of SRSF11 and SRRM1 in cancer associated splicing are less well known. SRSF11 is a regulator of neuronal microexon splicing ^59^, and has also been implicated in CRC metastasis, by regulating the AS of HSPA12A and resultant increased N-cadherin expression ^31^. Dysregulation of SRRM1 has been observed in several cancer contexts. In B-ALL, upregulation of SRRM1 is part of a high-risk gene signature and associated splicing program, that has a functional role in leukemia cell survival, proliferation and invasion ^60^. Elevated SRRM1 expression has also been shown to contribute to tumor cell invasion in hepatocellular carcinoma ^28^, as well as increased metastasis and aggressiveness in prostate cancer ^29^. Elevated plasma SRRM1 levels have been reported to distinguish high risk PCa patients from control individuals suggesting potential utility as a biomarker ^30^. In addition, SRRM1 was shown to promote expression of the oncogenic splice variant of CD44 in response to activated RAS signalling, suggesting that in addition to overexpression, oncogenic signaling may provide a mechanism to dysregulate SRRM1 activity ^38^. Related to this, we conducted the screen that identified SRRM1 as a regulator of NUMB E9 splicing in DLD1 cells which express activated RAS^G13D^. Coupled with the observation that activated RAS signaling also promotes NUMB E9 inclusion ^17^, these findings suggest a close link between SRRM1 regulated splicing events and activated RAS signalling in cancer.

In addition to promoting NUMB E9 inclusion, we identified an additional set of 24 SRRM1 and SRSF11 dependent cancer associated splicing events in genes that contribute to the cancer hallmarks. Indeed, the downregulation of SRRM1 resulted in reduced proliferation and colony formation of colon and lung cancer cell lines. We validated SRRM1 and SRSF11 regulation of specific splicing events including those in MNK2, TBX3, FOXM1, KAT5, ECT2 and DIAPH1, in DLD-1 and HCT116 colon cancer cells, as well as A549 lung cancer and MB-MDA468 breast cancer lines, supporting that notion that the observed splicing patterns may be common features of multiple tumour types. For example, among the SRRM1 and SRSF11 regulated ASEs that demonstrated oncogenic splicing patterns in COAD, LUAD and LUSC cancers was exon 14 in MKNK2, a serine threonine kinase that regulates translation through phosphorylation of eIF4E. MKNK2 undergoes alternative splicing of exon 14 to form functionally distinct isoforms MKNK2A and MKNK2B that act as a tumor-suppressor by phosphorylating p38 MAPK and inducing p38-mediated cell death, or activate eIF4E without triggering the p38 MAPK stress response respectively.

Whilst ECT2 and DIAPH1’s role in oncogenesis and metastasis has been previously established ^54,55,61,62^, the potential functional impacts of the alternative splicing events we identified has not been characterized. In our analysis of data derived from TCGA, ECT2 exhibits significantly elevated exon 4 inclusion in tumor vs normal samples from LUAD and LUSC patients, whilst DIAPH1 exhibits significantly increased exon 2 inclusion in tumor vs normal samples from COAD and LUSC patients. Both ECT2 exon 4 and DIAPH1 exon 2 sequences are within regions of the protein with predicted functions in subcellular localization and protein-protein interactions, and further studies are required to determine protein isoform specific activities.

Depletion of SRRM1 also resulted in significant changes in expression of a small group of genes including the reduced expression of WNT pathway-associated genes such as NOTUM ^63^, PRDX2 ^64,65^, DDB2 ^66^ and CCND1 ^67,68^ and elevated expression of the tumor-suppressive WNT-inhibitor CDKN2B^69,70^. The effects of SRRM1 on WNT related gene expression may be a consequence of alternative splicing events we identified within the C-clamp DNA binding region of TCF7L2 (TCF4), which forms a complex with β-catenin important for WNT target gene expression ^71–73^. Although we were unable to validate the specific splicing event regulated by SRRM1 due to the complexity of splicing in this region, these TCF7L2 isoforms are known to play differential roles sustaining oncogenic Wnt signaling ^73^.

We also found that SRRM1 regulates specific splicing events in the TBX3 and FOXM1 transcription factors, which both have roles in development, cancer, and are downstream effectors of WNT signaling ^74,75^. In agreement with altered TBX3, FOXM1 and TCF7L2 transcriptional activation function, SRRM1 loss caused reduced expression of potential target genes with functions in proliferative and invasive capacity, EMT, as well as resistance to apoptosis such as ALDH3A1^76^, CHI3L1^77^, FADS2 ^78^, METTL3 ^34^, PGK1 ^79^, and RFC4 ^80^. As well, SRRM1 knockdown reduced expression of oncogenic lncRNAs UCA1 ^81,82^ and DCST1-AS1 ^83^ and elevated expression of tumor-suppressive TP53AIP1 ^84^and ACTA2-AS1 ^85^.

A recurrent theme in oncogenic splicing is the reversion of alternative splicing programs back to embryonic patterns, resulting in the expression of protein isoforms that promote cellular phenotypes associated with tumor cells ^86^. Our study suggests that SRRM1 promotes a set of exon inclusion events including NUMB E9, TBX3 exon 2a, ECT2 exon 4, and PTBP1 exon 9 ^51,87,88^ which are both elevated in tumor tissue ^35^ and in early development ^89^. Indeed SRRM2, the main co-factor of SRRM1 has been shown to be vital in determining cell fate in early development and maintenance of cellular pluripotency ^90^. As well, TBX3 exon2a inclusion for example, has been shown to be important for maintenance of ES cell self renewal ^42^. Of note, multiple components of the core SAGA transcriptional co-activator complex, were identified as hits in our CRISPR screen, including SUPT7L, TAF5L, TAF6L, SUPT20H, and TADA1 ^91^, which regulates expression of RNA processing factors that activate an alternative splicing program during cellular reprogramming that regulates pluripotency ^92^.

Overall, this study has expanded the repertoire of known factors that regulate NUMB E9 splicing and further identifies the capacity of SRRM1 and SRSF11 to promote oncogenesis by coordinately regulating a program of oncogenic splicing events. This work also highlights SRRM1 as a potential therapeutic target, by linking its function to proliferation, EMT, apoptosis resistance, stemness and the regulation of WNT signaling.

## Materials & Methods

### Construction of human Numb exon 9 dual fluorescent splicing reporters

A dual fluorescent vector was constructed by cloning EGFP from a N1-EGFP vector into the N-terminal of a N2-mCherry vector such that EGFP and mCherry proteins are in alternate reading frames. To generate the splicing reporter, we utilized the human Numb genomic sequence chr14: 72778081-72952456 (NCBI build 36.1/hg18) obtained in a pBACCe3.6 vector (Empire Genomics, Buffalo, NY, USA; cat. #RP11-76J16) as previously described by our lab ^17^. The human genomic region encompassing Numb exons 8/9, the initial fragment of exon 10, and introns 8-9/9-10 (5302 nucleotides total) were amplified by PCR and subcloned into the dual fluorescent vector.

To allow for expression of EGFP or mCherry fusions depending on the inclusion or skipping of exon 9 (E9), a frameshift mutation inserting two additional nucleotides was introduced into Numb E9 using site directed mutagenesis (TAGCACCCGTA**<u>tt</u>**GCAATGCCTGT; inserted nucleotides are bolded and underlined). To ensure that the frameshift mutation did not alter the splicing of the E9 reporter, we examined the effect of insertion of one (E9+1) or two (E9+2) additional nucleotides on E9 splicing when inserted into a 2HA-Numb E9 reporter ^17^ (Supp Fig 1 A, B). Insertion of two nucleotides (E9+2) did not alter Numb reporter splicing relative to the non-mutagenized version of the reporter, and was thus cloned into a pcDNA5/FRT/TRE-based plasmid compatible for insertion in Flp-In T-REx cells (Thermofisher: Cat: K650001). The pcDNA5 vectors were then used to generate stable dual fluorescent Flp-In HEK293 and DLD1 Numb E9 reporter cell lines.

### Generation of stable Flp-In-DLD1/Flp-In-HEK reporter cell lines

Tetracycline/doxycyline-inducible stable cell lines were generated by transfecting 1 μg of pcDNA5/FRT/TRE-based Numb E9 reporter plasmid and 500 ng of plasmid encoding pOG44 recombinase in HEK293 Flp-In or DLD1 Flp-In lines using Lipofectamine 3000 (Life Technologies) according to the manufacturer’s instructions. Cell lines with stably integrated constructs were selected for and maintained with 15 μg/mL blasticidin and 200 μg/mL hygromycin. Transgene expression was induced by addition of 10 μg/mL tetracycline.

### Cell culture

Flp-In DLD1 colorectal adenocarcinoma cells were generously donated by the Gingras Lab. DLD1 cell lines were grown in RPMI 1640x medium (Wisent Bioproducts: Cat: 350-045-CL). HCT116 colorectal carcinoma cell line was grown in McCoy’s 5A medium (WISENT, Cat: 317-010-CL). A549 and HEK293 cell lines were grown in DMEM (WISENT, Cat: 319-005-CL). MDA-MB-468 human breast cancer cell line was grown in Leibovitz L-15 medium (Gibco, Cat: 21083-027). All media were supplemented with 10% fetal bovine serum (FBS) and 1% penicillin/streptomycin. Cell lines were authenticated based on STR profiling using services from The Centre for Applied Genomics in SickKids.

### DNA transfection

Cells were grown to 70-80% confluence in 6-well plates (Nunc) and were transfected with 1 μg of plasmid DNA 18-20 hours after seeding. Cells were transfected with plasmids expressing known Numb E9 splicing regulators previously established by our group ^17^ (SRSF1, PTBP1, constitutive active form of MEKK, MEKK-EE) or an empty control vector (pcDNA3.1+) using Lipofectamine 2000 (Invitrogen) and Opti-MEM reduced serum medium (Life Technologies) according to the manufacturer’s directions. Cells were maintained in serum-supplemented medium during the transfection process. 48-72 hours post-transfection, cells were lysed in the appropriate lysis buffer for either RNA or protein extraction and validation of Numb reporter splicing via immunoblots/RT-qPCR.

### Immunoblots

Cells were washed twice with PBS and lysed with 1X Laemmli Sample Buffer (63mM TrisHCl pH6.8, 2% SDS, 10% Glycerol). Cell lysates were collected into microfuge tubes, boiled for ten minutes, mixed for 10 minutes and centrifuged for 10 minutes at maximum speed. A BCA assay (Pierce) was used to determine the protein concentration in each lysate. A mixture of 2-mercaptoethanol (final concentration 5%) and bromophenol blue (final concentration 0.001%) was added to each lysate and 50 μg of protein were loaded onto 8-12 % gels for SDS-PAGE. Proteins were then transferred to PVDF membranes by wet transfer and blocked with 1% fish gelatin (Millipore Sigma, #G7765) in TBS. Membranes were incubated in primary antibody (listed below) overnight. The following day, membranes were washed three times in TBST for 10 minutes each, then incubated with HRP conjugated secondary antibodies for 1 hour followed by another set of TBST washes before development with ECL substrates. BioRad Gel Doc Imager was used to capture the signal after addition of ECL substrates. Membranes were stained with Coomassie blue stain for quantification of total protein expression. BioRad’s Image Lab software was used to quantify band signals.

- Rabbit anti-EGFP (Invitrogen, Ref: A-11122) at 1:1000
- Rabbit anti-mCherry (Rockland, Ref: 600-401-p16) at 1:1000
- Rabbit anti-Gaussia luciferase (Thermofisher, Cat: PA1-181) at 1:1000
- Rabbit anti-Red Firefly luciferase (Thermofisher, Cat: PA1-179) at 1:1000
- Rabbit anti-SRRM1 (Invitrogen, Ref: PA5-62100) at 1:1000
- Mouse anti-Flag M2 (Sigma-Aldrich, Ref: F1804-200UG) at 1:1000
- Rabbit anti-phospho-p38 (Cell Signaling Technology, Ref: D3F9) at 1:1000
- Mouse anti-p38 (Santa Cruz, Ref: sc-7972) at 1:1000
- Mouse anti-GAPDH (Invitrogen, Ref: MA5-15738) at 1:10000

### Splicing factor KO FACs test screen

Five lentiCRISPRv2 vectors expressing guides targeting mCherry, SRSF1, SRSF3 and PTBP1 were obtained from Sigma Aldrich (gRNA sequence shown in Table 1). The lentiCRISPRv2 vector targeting AAVS1 (adeno-associated virus integration site 1) used as a negative control was generously provided by the Angers lab. The lentiCRISPRv2 vectors were used to examine whether knockdown of known Numb E9 splicing regulators was capable of influencing reporter EGFP/mCherry expression as examined by FACS. DLD1 Flp-In cells expressing the Numb E9 splicing reporter were transduced at an MOI of 0.3 in the presence of 8 mg/mL polybrene. 24 hours after infection, the medium was replaced with fresh medium containing puromycin (2.0 mg/mL) and cells were incubated for an additional 72 hours before passaging. 12 days post infection, cells were harvested and fixed in ice cold PBS buffer containing 1% PFA for 30 minutes and then washed twice with PBS. Cells were passed through a nylon mesh with a pore size of 40 mm to eliminate large aggregates and then sorted based on the relative EGFP:mCherry expression as indicated in (Fig 1C) using the Flow Cytometry core facility at the Peter Gilgan Centre for Research and Learning. Gates were generated to capture the top 5% mCherry high:EGFP low and EGFP high:mCherry low populations using the negative control sg-AAVS1 cells and were maintained for all the other targeted genes to examine shifts in the dual fluorescent population in response to their KO.

### Virus library production and MOI determination

For virus library production, 9 million HEK293T cells were seeded on 15 cm plates and transfected 24 hours later with a mix of 14 μg lentiviral pLCKO vector containing the library, 16μg packaging vector psPAX2, 1.56μg envelope vector pMD2.G, 93.6 μL X-treme Gene transfection reagent (Roche) and 1.4 mL Opti-MEM medium (Life Technologies). 24 hours post-transfection the medium was changed to serum-free, high BSA growth medium (DMEM, 1.1 g/100mL BSA, 1% penicillin/streptomycin). The virus-containing medium was harvested 48 and 72 hours after transfection, centrifuged at 1,500 rpm for 5 minutes and then frozen. DLD1 dual-fluorescent reporter cells were infected with a titration of the lentiviral gRNA library along with polybrene (8 μg/mL). After 24 hours the medium was replaced with medium containing puromycin (2 μg/mL) and cells were incubated for an additional 48 hours. Multiplicity of infection (MOI) of the titrated virus was determined 72 hours post-infection by comparing the percent survival of infected cells to a non-infection control.

### Genome-wide CRISPR-Cas9 screen

100 million DLD1 Flp-In cells expressing the dual fluorescence splicing reporter were infected with the lentiviral TKOv3 library (70,948 sgRNAs) at an MOI of 0.3 in the presence of 8 mg/mL polybrene to achieve a library coverage of ∼300 cells/gRNA. 24 hours after infection, the medium was replaced with fresh medium containing Puromycin (2.0 mg/mL) and cells were incubated for an additional 72 before collecting a T0 unsorted cell population for technical analysis of the CRISPR screen. After selection, cells were split such that each sgRNA would be represented by an average of 300 cells in the population and passaged every three days. For cell sorting, 30 million cells were seeded and the following day 1μg/mL of tetracycline was added to induce dual fluorescent reporter expression. 24 hours later the cells were harvested and fixed with 1% PFA in ice cold PBS for 30 minutes and resuspended in sorting buffer (Hanks Balanced Salt Solution, 25mM HEPES pH 7.0, 1mM EDTA, 1% albumin) at a concentration of 10 million cells per millilitre. Cells were passed through a nylon mesh filter with a pore size of 40 mm to eliminate large aggregates. Filtered cells were sorted based on the relative EGFP:mCherry expression using Flow Cytometry core facilities at the Peter Gilgan Centre for Research and Learning using a Beckman Coulter MoFlo Astrios Cell Sorter. For each replicate, 900,000-1,100,00 cells with the highest or lowest 5% EGFP:mCherry ratio were collected. Additionally, 30 million cells were collected prior to sorting as a reference point for comparing enrichment in the sorted populations. The same procedure was followed for sorting of cells with the 20% highest or lowest EGFP:mCherry ratio.

Genomic DNA was extracted from the cell pellets of unsorted samples using the QIAamp Blood Maxi Kit (QIAGEN) while gDNA from sorted populations was purified with the Qiagen Midi Kit as per the manufacturer’s recommendations. gDNA was precipitated using ethanol and sodium chloride then resuspended in sterile water. gRNA inserts were amplified via PCR and prepared for sequencing using primers harboring Illumina TruSeq adapters with i5 and i7 barcodes. The resulting libraries were sequenced on an Illumina HiSeq2500 at the Donnelly Sequencing Centre. Each read was completed with standard primers for dual indexing with Rapid Run V1 reagents. The first 20 cycles of sequencing were “dark cycles”, or base additions without imaging. The actual 26 base pair reads began after the dark cycles and contained two index reads with the i7 sequence read first, and the i5 sequence read second.

### CRISPR screen analysis

Sequencing reads were mapped to TKOv3 library sgRNAs using the count function in the MAGeCK software (v0.5.8). Gene and sgRNA level enrichment in the mCherry high:EGFP low and EGFP high:mCherry low cell populations were calculated using the test function in MAGeCK in comparison to unsorted cells with control gRNAs targeting LacZ, EGFP and luciferase excluded during analysis. Genes over-represented at an FDR < 0.2, and a fold enrichment of at least 1.5x (Log fold enrichment > 0.585) with either gating regimen (5% or 20% highest or lowest mCherry:EGFP ratio) were considered as hits.

Gene ontology biological processes, molecular function and cellular component term enrichment analysis of all unique hits from all sorted populations (FDR < 0.2, LFC > 0.585 with gene expression > 1.0 TPM in DLD1 cells according to Depmap CCLE https://depmap.org/portal/ccle/) were uploaded to the online Database for Annotation, Visualization, and Integrated Discovery (DAVID, https://david.ncifcrf.gov/home.jsp), and clustered using an EASE of 1.0 and medium stringency. STRING interaction network of genes shown in (Fig 2D) identifying a potential regulatory splicing complex operating on Numb E9 alternative splicing was conducted using string database (https://string-db.org/). Only curated databases and experimentally determined evidence were considered. Nodes were colored based on biological process and molecular function go terms related to mRNA splicing, via spliceosome, RNA splicing, RNA binding and mRNA processing.

### siRNA transfections

DLD1, HCT116, A549, MDA-MB-468 and HEK293 cell lines were transfected with 25 nM of On-TARGETplus Smartpool siRNA (a mixture of four siRNA provided as a single reagent; Horizon Discovery) using Dharmafect 2 transfection reagent (Horizon Discovery) and Opti-MEM (Life Technologies) as per the manufacturer’s instructions. A non-targeting siRNA pool was used as control. Transfected cells were harvested 48 hours or 72 hours post transfection for RNA collection using Qiagen’s RNeasy RNA extraction kit. RNA samples were used for RT-qPCR and RT-PCR experiments.

### RT-qPCR

1μg of each RNA sample was reverse transcribed using the High-capacity cDNA Reverse Transcription kit (Applied Biosystems) and a prepared mixture of random oligomers and oligo-dT primers following the manufacturer’s instructions. cDNA samples were diluted as required and 2 μL of dilute cDNA was added to 18 μL of the appropriate real-time reaction master mix in a 96-well clear PCR plate. Real-time reaction master mixes were prepared with PowerUp SYBR Green Master Mix (Applied Biosystems) with 12.5 μM of each forward and reverse primer according to the manufacturer’s protocol. Reactions were set up in triplicate. RT-qPCR reactions were run on an Applied Biosystems Quantstudio3 thermal cycler, using cycling conditions described by the manufacturer, and an annealing temperature of 60°C. Reactions were normalized to GAPDH expression. A list of RT-qPCR primers are described in (Table 2).

**Table 2:**
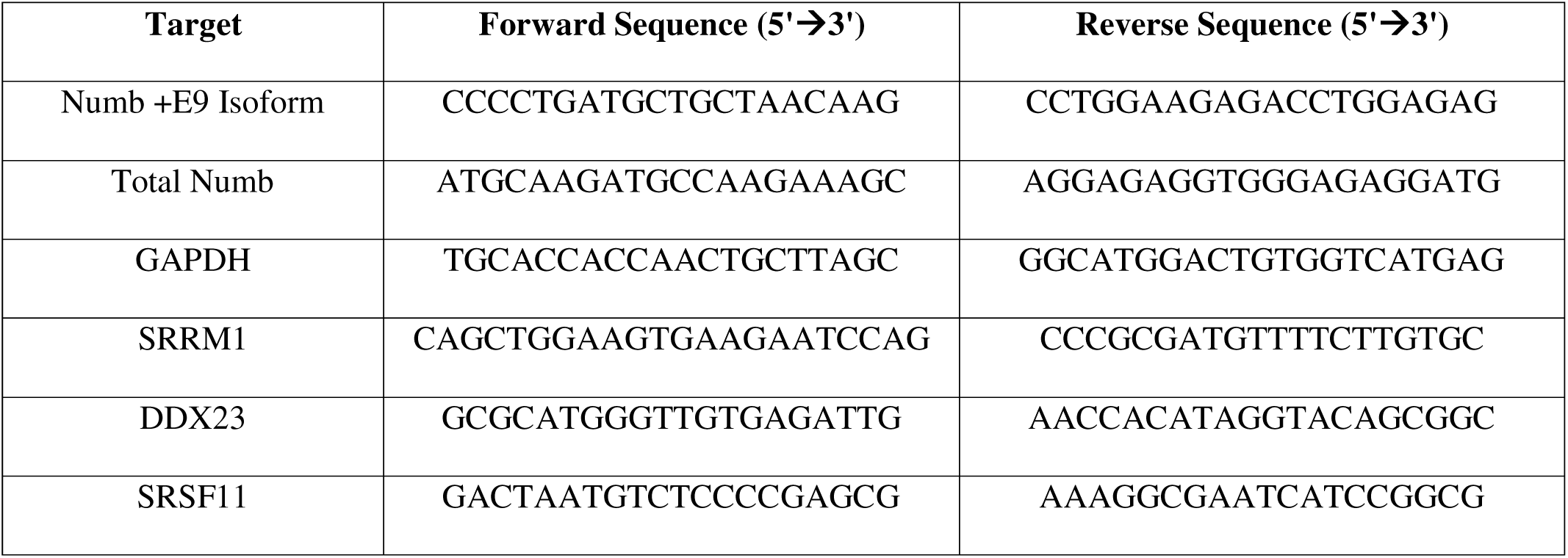
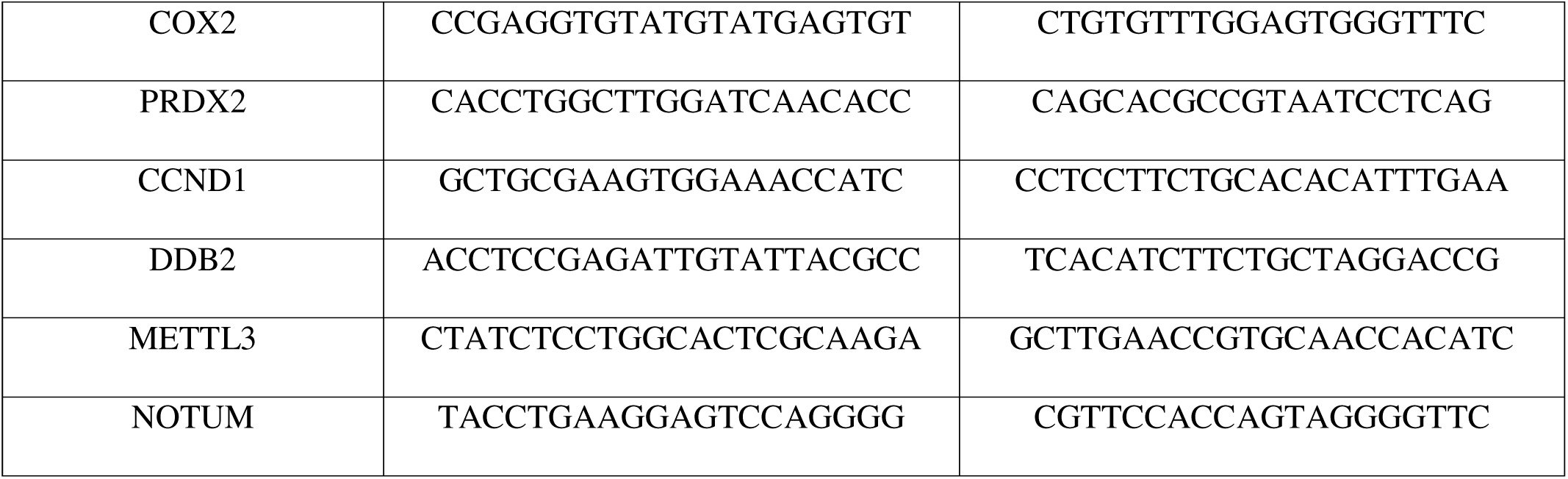
List of primers used for quantitative PCR.

### Semi-quantitative PCR

1 μg of RNA was reverse-transcribed using the High-capacity cDNA Reverse Transcription kit (Applied Biosystems) and a prepared mixture of random oligomers and oligo-dT primers following the manufacturer’s instructions. 1/20^th^ of the cDNA reaction volume was used for PCR amplification with 12.5 μM of isoform-specific primers and 0.25 μL of Taq DNA polymerase (Roche) in a final reaction volume of 50 μL. PCR reactions were run on a BioRad thermocycler and had an initial denaturing step at 94°C for 2 minutes, followed by multiple cycles of denaturing, annealing and 72°C elongation, before a final elongation step for seven minutes at 72°C. PCR products were then resolved on 1-3% agarose gels pre-stained with RedSafe dye (Chembio). A BioRad gel doc was used to capture images, and BioRad’s Image Lab software was used to quantify band signal intensity. A list of primers used for semi-quantitative PCR can be found in (Table 3).

**Table 3:**
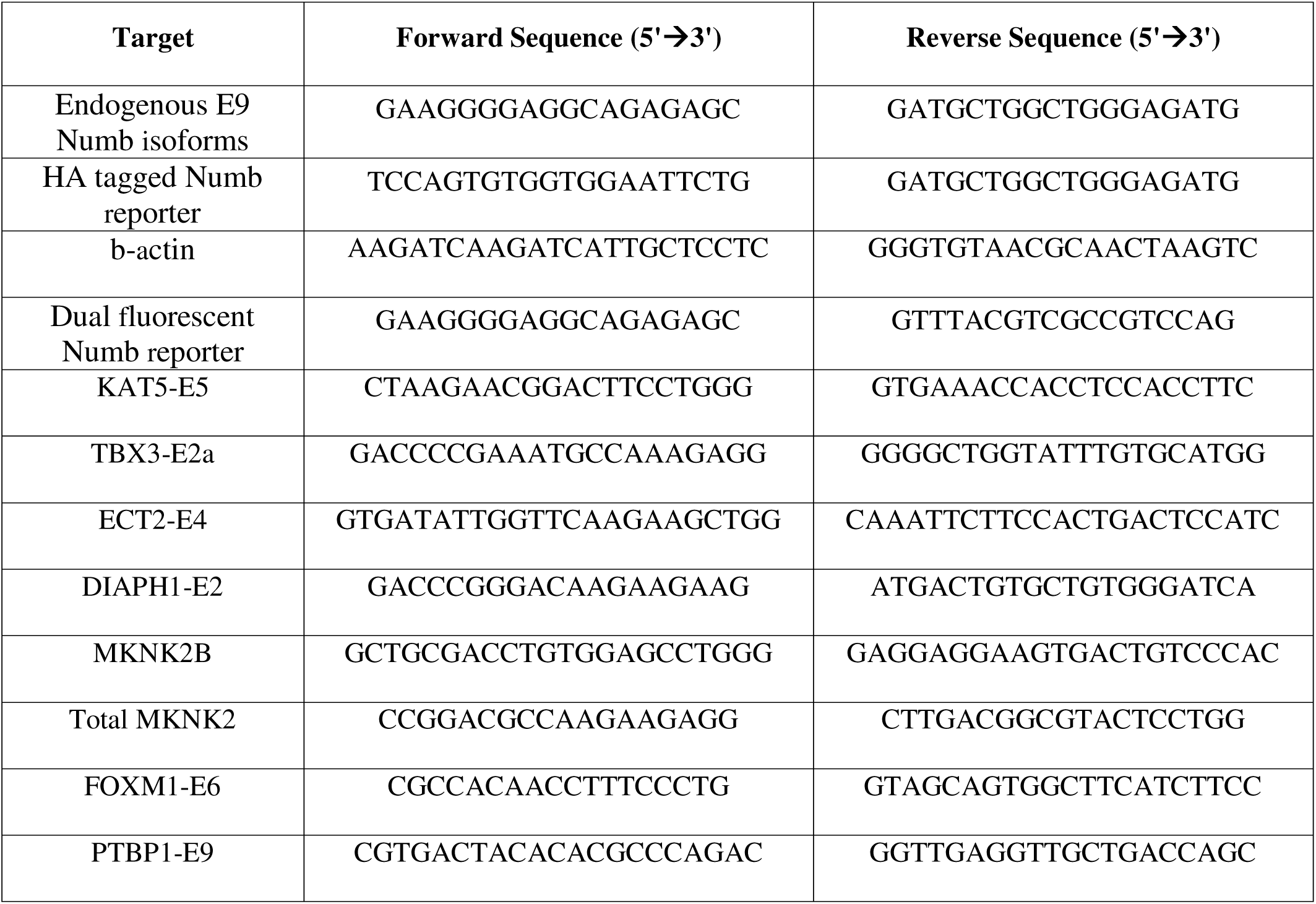
List of primers used for semi-quantitative PCR.

### Generation of flag-tagged SRRM1 and SRSF11 DLD1 and HEK293 cell lines

1 μg of pcDNA5/FRT/TRE-based plasmid encoding FLAG-tagged SRRM1 or SRSF11 was co-transfected with 500 ng of pOG44 recombinase plasmid in DLD1 or HEK293 Flp-In lines with Lipofectamine 2000 (Life Technologies) according to the manufacturer’s protocol. Parental Flp-In lines were maintained with media containing 15 µg/mL blasticidin and 100 µg/mL zeocin. To select for cell lines with stably integrated constructs, cells were grown in media containing 15 μg/mL blasticidin and 200 μg/mL hygromycin. Transgene expression was activated by supplementing media with 10 µg/mL of tetracycline.

### FLAG Affinity Purification coupled with Mass Spectrometry (AP-MS) sample preparation

Flag-tagged SRRM1 or SRSF11 DLD1 or HEK293 cell lines were seeded onto two 15 cm plates and induced for 24 hours with 1 μg/mL tetracycline. Cell pellets were collected and lysed in ice cold TAP lysis buffer containing 50 mM HEPES-NaOH (pH 8.0), 100 mM NaCl, 2 mM EDTA, and 10% glycerol with freshly added 0.1% NP-40, 1 mM DTT, 1 mM PMSF, and a single COMPLETE protease inhibitor tablet (Roche Applied Science) at a 1:6 pellet weight-to-volume ratio. Resuspended pellets were frozen and thawed by placing the pellets on dry ice for five minutes and thawed in a 37°C water bath until a small ice pellet remained. Samples were quickly moved back onto ice and sonicated with three 10 second bursts with a 2 second rest at an amplitude of 35%. To solubilize chromatin and reduce the detection of interactions mediated by RNA or DNA, 1 μL of benzonase nuclease (Sigma-Aldrich, E8263, 250U) was added to each lysate and incubated for 30 minutes with rotation at 4°C. Lysates were cleared by centrifugation at 20,000 rcf for 20 minutes at 4°C then transferred to tubes containing 25 μL of 50% magnetic anti-FLAG M2 beads slurry (Sigma-Aldrich, M8823) pre-washed with lysis buffer. FLAG immunoprecipitation was performed for 3 hours at 4°C with rotation. After incubation, beads were pelleted by centrifugation at 1,000 rpm for 1 minute and magnetized to aspirate the unbound lysate. The beads were then de-magnetized and washed with 1 mL of lysis buffer and the total volume (with beads) were transferred to a new tube. Beads were washed once more with 1 mL of lysis buffer then washed with 50 mM ammonium bicarbonate (ABC) at pH 8. All wash steps were performed on ice using cold lysis buffer and ABC.

All sample preparation, LC-MS/MS data collection, and MS data searches were carried out at the SPARC BioCentre at The Hospital for Sick Children (Toronto, ON, Canada). Samples were reduced with DTT (10 mM, 60°C, 1 hour), alkylated with iodoacetamide (20 mM, room temperature, 45 minutes, dark), and digested overnight at 37°C with trypsin (2 μg; Pierce). Peptides were dried by vacuum centrifugation, desalted on C18 ziptips (Millipore) using a DigestPro MSi (Intavis Bioanalytical Instruments), and dried again by vacuum centrifugation before resuspension in Buffer A (2% acetonitrile, 0.1% formic acid). Samples were analyzed by liquid chromatography tandem mass spectrometry (LC-MS/MS) using an EASY-nanoLC 1200 system with a 1 h analysis and an Orbitrap Fusion™ Lumos™ Tribrid™ Mass Spectrometer (Thermo Fisher Scientific). The LC portion of the analysis consisted of a 18 minute linear gradient running 3-20% of Buffer A to Buffer B (0.1% FA, 80% acetonitrile), followed by a 31 minute linear gradient running 20-35% of Buffer A to Buffer B, a 2 minute ramp to 100% Buffer B and 9 minute hold at 100% Buffer B, all at a flow rate of 250 nL/minute. Samples were loaded into a 75 μm x 2 cm Acclaim PepMap 100 Pre-column followed by a 75 µm x 50 cm PepMax RSLC EASY-Spray analytical column filled with 2 µM C_18_ beads (Thermo Fisher Scientific). MS1 acquisition resolution was set to 120000 with an automatic gain control (AGC) target value of 4 x 10^5^ and maximum ion injection time (IT) of 50 ms for a scan range of *m/z* 375-1500, with dynamic exclusion set to 10 s. Isolation for MS2 scans was performed in the quadrupole with an isolation window of *m/z* 0.7. MS2 scans were performed in the ion trap with maximum ion IT of 10 ms, AGC target value of 1 x 10^4^, and higher-energy collisional dissociation (HCD) activation with an NCE of 30.

### Mass spectrometry data analysis

MS raw files were analyzed using PEAKS Studio software (Bioinformatics Solutions Inc.) and Proteome Discoverer (version 2.5.0.400) and fragment lists searched against the mouse UniProt Reference database (Uniprot_UP000000589 downloaded Sept. 15, 2020; 21966 entries). For both search algorithms, the parent and fragment mass tolerances were set to 10 ppm and 0.6 Da respectively, and only complete tryptic peptides with a maximum of three missed cleavages were accepted. Carbamidomethylation of cysteine was specified as a fixed modification.

Deamidation of asparagine and glutamine, oxidation of methionine, acetylation of the protein N-terminus, and phosphorylation of serine, threonine, and tyrosine residues were specified as variable modifications. Membership in spliceosomal sub-complexes of prey proteins identified in AP-MS experiments was analyzed using human protein annotations in the Spliceosome Database (^93^). In (Fig 3A, Supp Fig 4A), only prey with a saint score > 0.95 for at least one of the baits are shown. The network map was generated using Cytoscape. Gene ontology biological processes term enrichment analysis of all unique SRRM1 or SRSF11 protein interactors (BFDR < 0.05) in DLD1 cells or (Saint score > 0.95) in HEK293 were uploaded to the online Database for Annotation, Visualization, and Integrated Discovery (DAVID, https://david.ncifcrf.gov/home.jsp), and clustered using an EASE of 1.0 and medium stringency. Data are available for download at MassIVE: MSV000100469

### Bulk RNA-seq data analysis

Raw reads were mapped to the human genome using the hg19 reference genome with the STAR aligner (version 2.7.11a) ^94^. PCR duplicates and low-mapping quality reads were removed using Picard tools and SAMtools. Differential expression analysis was performed using the R package DESeq2 (v.1.42) ^95^. Only genes with a FDR adjusted P value < 0.05 and foldchange > 2 were considered statistically significant and graphed as a volcano plot using ggplot2 packages on R software. Differential splicing analysis was performed using rMATS-turbo (version 4.2.0) ^96^. Splicing events supported by at least 15 unique reads with a ΔPSI > 10% and an FDR of < 0.05 were considered statistically significant. Statistical analysis was performed and plots were generated using R environment version 4.3.1. Sashimi plots were generated using ggsashimi.py script ^97^ with the parameter –min-coverage 5 and –aggr mean. Motif enrichment analysis was performed using rMAPS ^98^. Significantly differently expressed genes with SRRM1 knockdown were uploaded to ChEA3 online tool (https://maayanlab.cloud/chea3/) ^57^ to predict transcription factor regulation. The transcription factors predicted to regulate these genes were cross referenced to the list of differentially spliced genes to identify specific transcription factors that may mediate the change in gene expression observed with SRRM1 knockdown (supplementary data).

Gene ontology biological processes term enrichment analysis of all unique genes that had alternatively spliced exons (ASEs) regulated by SRRM1 or SRSF11 were uploaded to the online Database for Annotation, Visualization, and Integrated Discovery (DAVID, https://david.ncifcrf.gov/home.jsp) and were clustered using an EASE of 1.0 and medium stringency. Gene set enrichment analysis (GSEA) of all differentially expressed genes in response to SRRM1 knockdown was conducted using GSEA software from the Broad institute using a pre-ranked list (SIGN(logFC)*-log10(pvalue)), maximum term size (500), minimum term size (15) and 2000 permutations using MSigDB gmt file (h.all.v2024.1.Hs.symbols.gmt) obtained from (https://www.gsea-msigdb.org/gsea/index.jsp) ^99,100^. RNAseq data are available at DOI: 10.5281/zenodo.18154912.

### TCGA alternative splicing analysis

Our study utilized alternative splicing analysis that was conducted by ^35^ on RNA-sequencing (RNA-seq) samples from TCGA database. BRCA, COAD, LUAD and LUSC PSI values were extracted from Kahles et al., supplementary tables for all alternative splicing events that involved genes identified to be differentially spliced from our own DLD1 bulk RNA-seq analysis. A two-sided unpaired wilcoxon test of the difference of the PSI value between tumor and normal samples in BRCA, COAD, LUAD and LUSC was conducted. Specific alternative splicing events that demonstrated significant differences between tumor and normal samples (p-value < 0.05) were matched to the specific alternative splicing events from our bulk RNA-seq data analysis using the specific coordinates of the upstream, spliced and downstream exons using the GRCh73/hg19 human genome assembly. These steps were conducted using R environment version 4.4.1. GraphPad Prism software was used to graph the difference between PSI in tumor and normal samples (BRCA/COAD/LUAD/LUSC PSIΔ) versus the difference between PSI in the NTC and SRRM1/SRSF11 knockdown conditions (SRRM1si/SRSF11si PSIΔ) to determine similarity and differences between TCGA PSI changes and DLD1 RNA-seq PSI changes (Supp Fig 7/9).

For TCGA PSI analysis of MKNK2 E14a, ECT2 E4, DIAPH1 E2, KAT5 E5 and TBX3 E2a (Supp Fig 9A), the hg19 chromosomal coordinates of each splicing event used for extraction is described in (Table 4). PSI values and sample barcodes were imported into R Studio version 4.41. Sample barcodes were broken down to identify the cancer (i.e., BRCA, COAD, LUAD etc.) and sample type (i.e., normal solid tissue, primary solid tumor, metastatic). The PSI values for each splicing event were compared between primary solid tumor samples and normal solid tissue samples within each cancer type. Only cancer types with five or more paired normal tissue samples were included in the analysis. A Wilcoxon rank-sum test was performed to determine if the difference in PSI for each splicing event between normal and tumor samples were statistically significant for each cancer type. Figures were created using GraphPad Prism.

**Table 4:**
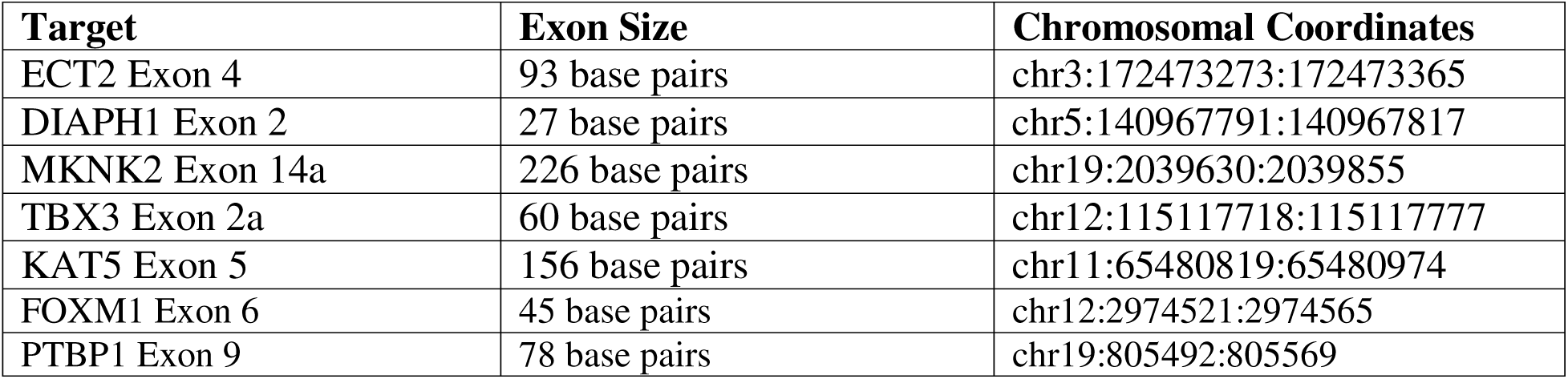
List of chromosomal coordinates used in TCGA PSI analysis.

### Wound Healing Migration and Invasion Assays

For HCT116 wound healing assay, 80,000 cells (shNTC, shSRRM1 611 and shSRRM1 960) were seeded in 96-well incucyte imagelock plates (Sartorius) coated with 100 ug/mL of matrigel (Corning BD Cat. No. 354234). HCT116 cells were seeded in normal media. For A549 wound healing assays, cells were induced with 10 ug/mL of tetracycline for 72 hours prior to seeding. On day of seeding, 70,000 cells (shNTC, shSRRM1 611, and shSRRM1 960) were seeded in 96-well incucyte imagelock plates coated with matrigel. A549 cells were seeded in media supplemented with tetracycline. After 24 hours, the cell monolayer (at 90–100% of confluence) was wounded by scraping with the Incucyte® Wound Maker 96-Tool (Sartorius).

Cells were washed with medium to remove floating cells and replaced with medium supplemented with 10% FBS, 1% penicillin/streptomycin and 10 μg/ml tetracycline. Relative wound density was imaged and measured with an Incucyte® Live-Cell imager (Sartorius) every 4 hours post-wounding. The wound width was evaluated in a minimum of four wells using the Incucyte® Cell Migration Scratch Wound Analysis software. For the invasion assay, Matrigel (4 mg/mL, Corning BD Cat. No. 354234) was added after wounding and washing following Sartorius Incucyte guidelines. The mean of three independent experiments from shNTC, shSRRM1 611 and shSRRM1 960 cell lines were analyzed using GraphPad Prism by conducting a two-way ANOVA comparing the NTC to SRRM1 knockdown condition to determine the statistical significance of SRRM1 knockdown on cell migration and invasion.

### Proliferation assays

For the HCT116 proliferation assay, 2,500 cells (shNTC, shSRRM1 611 shSRRM1 and 960 HCT116) were seeded in 96-well incucyte imagelock plates (Sartorius). After 24 hours, cells (at ∼10% of confluence) had their medium replaced with fresh medium supplemented with 10% FBS, 1% penicillin/streptomycin and 10 μg/mL tetracycline. For the A549 proliferation assay, 2,500 cells (shNTC, shSRRM1 611, shSRRM1 960) were seeded in 96-well incucyte imagelock plates in medium supplemented with tetracycline. Phase area confluence was imaged and measured with an Incucyte® Live-Cell imager (Sartorius) every 6 hours. Cell confluence was evaluated in a minimum of eight wells using the Incucyte® AI confluence and basic fluorescence analysis software following Sartorius Incucyte guidelines. The mean of three independent experiments from shNTC, shSRRM1 611 and shSRRM1 960 cell lines were analyzed using GraphPad Prism by conducting a two-way ANOVA comparing the NTC to SRRM1 knockdown condition to determine the statistical significance of SRRM1 knockdown on cell proliferation.

### Colony formation assays

675, 1250 and 2500 cells (shNTC, shSRRM1 611, shSRRM1 960 HCT116) were seeded in 6-well plates and cultured in standard conditions. Fresh medium was replaced every week. After 14 days, cells were fixed and stained with crystal violet (0.1%w/v). Plates were scanned to generate representative images. The crystal violet stain was then solubilized using 1% SDS and the signal was acquired by measuring absorbance at 590 nm. Statistical significance was determined using paired t-test between shNTC and shSRRM1 biological triplicates.

## Acknowledgements

The authors thank the following: SPARC molecular analysis, the Sickkids Flow cytometry facility, Donnelly Sequencing Centre and Thomas Gonatopoulos-Pournatzis and Benjamen Blencowe for technical advice, Anne Claude Gingras for providing Flp-In DLD1 cell line. This work was supported by funding from the Canadian Institutes of Health Research (Project Grant FRN 106507) and the Canadian Cancer Society (Challenge Grant - 707475) and Natural Sciences and Engineering Council of Canada (RGPIN-2019-06485) to CJM. LV is a recipient of a Canada Graduate Scholarship – Masters. HF is a recipient of an Ontario Graduate Scholarship. CC and RB received an USRA awards from NSERC.

## Author Contributions

KO performed experiments, acquired and analyzed data, drafted and critically reviewed the manuscript. LV, HF, JL, CDS, CC, RB AZ performed experiments, acquired and analyzed data. GM, SA analyzed and interpreted data. OS analyzed and interpreted data and helped draft the manuscript. CJM supervised KO, LV, HF, CC, RB, AZ, analyzed and interpreted data, drafted and revised the manuscript.

