## Supplementary Figures for "SRRM1 coordinates an alternative splicing program that promotes expression of oncogenic protein isoforms"

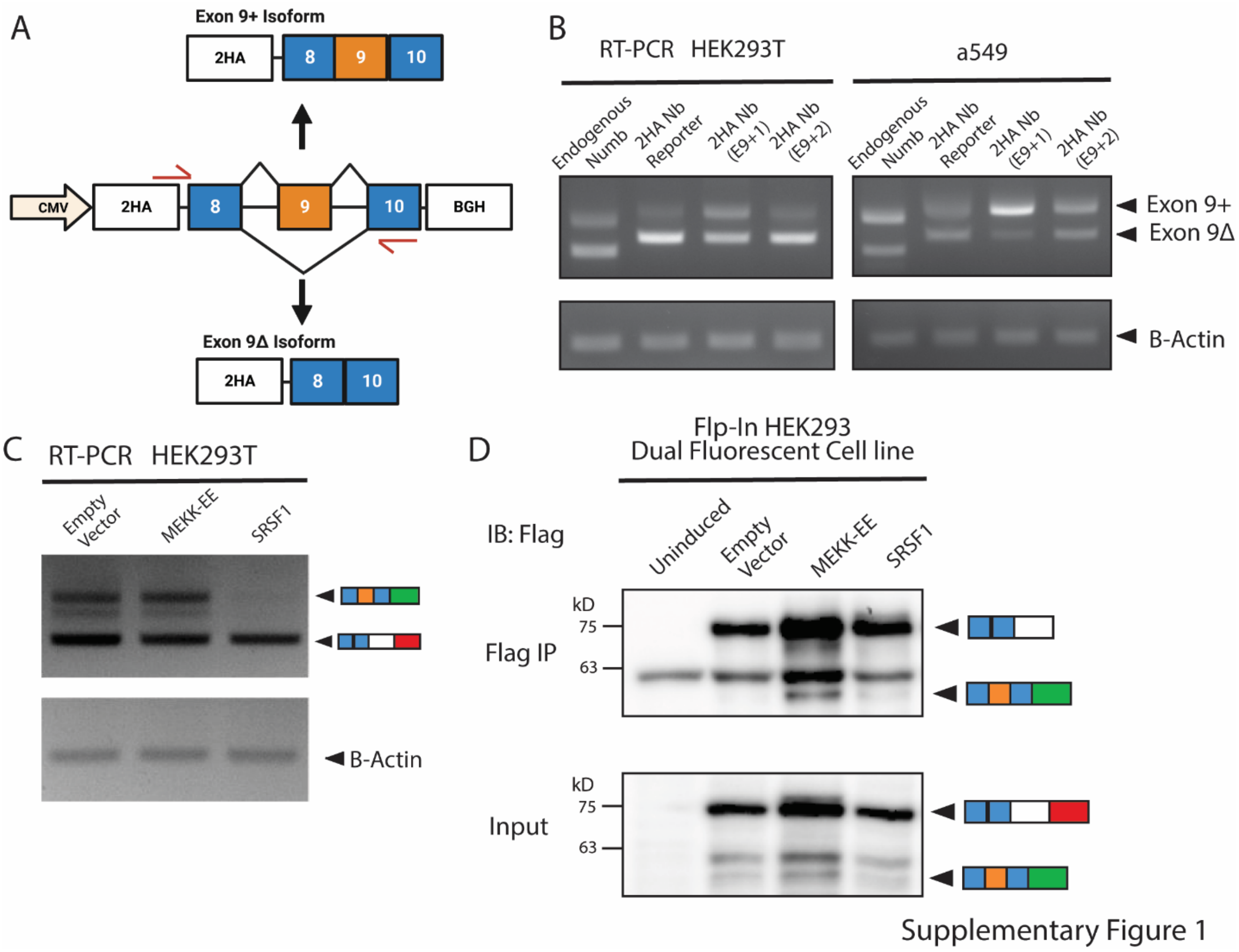


**Supplementary Figure 1: Generation of tetracycline-inducible dual fluorescent Numb E9 splicing reporter cell lines.**

**(A)** Schematic of the HA-tagged Numb E9 splicing reporter originally described in (Rajendran 2016). The backbone is pcDNA3.1+ vector with an amino-terminal double-HA tag, CMV promoter and a BGH and polyadenylation signal. A fragment of human Numb genomic DNA containing the entire region from exons 8 through 10 was cloned into the vector. (chr14; 72811671-72818966, hg 18). The expression of the reporter produces 2 transcript variants of Numb. The primers sites used to target the reporter transcripts by RT-PCR are indicated by red arrows on the reporter. This reporter was used to test the effect of inserting frameshift mutations within exon 9 on reporter splicing

**(B)** Expression of different variants of the 2HA Numb splicing reporter in HEK293T cells and a549 cells. Total RNA was collected from cells transfected and cDNA was subject to PCR using primers shown in a **(A)** as well as primers targeting endogenous exon 8 and 9, comparing the expression of exon 9 in two modified versions of the reporter, which were mutagenized to add either a single base pair (E9+1) or two base pairs (E9+2) in the middle of exon 9, to demonstrate the effect of mutagenesis on reporter splicing relative to endogenous Numb.

**(C)** Co-expression of the dual fluorescent splicing reporter in a549 and HEK293T cells with either a control empty vector (pcDNA3.1+) or constitutively active MEKK (MEKK-EE) or the splicing factor SRSF1. Total RNA was collected from transfected cells and cDNA was subject to RT-PCR using primers targeting exon 8 and EGFP, comparing expression of exon 9 isoforms.

**(D)** Representative immunoblots of exon 9 reporter isoforms in Flp-in HEK293 dual fluorescent reporter cell line lysates, which underwent anti-flag immunoprecipitation and probed with anti-flag antibodies. Lanes represent uninduced control cells, or induced cells transfected with empty vector, MEKK-EE or SRSF1

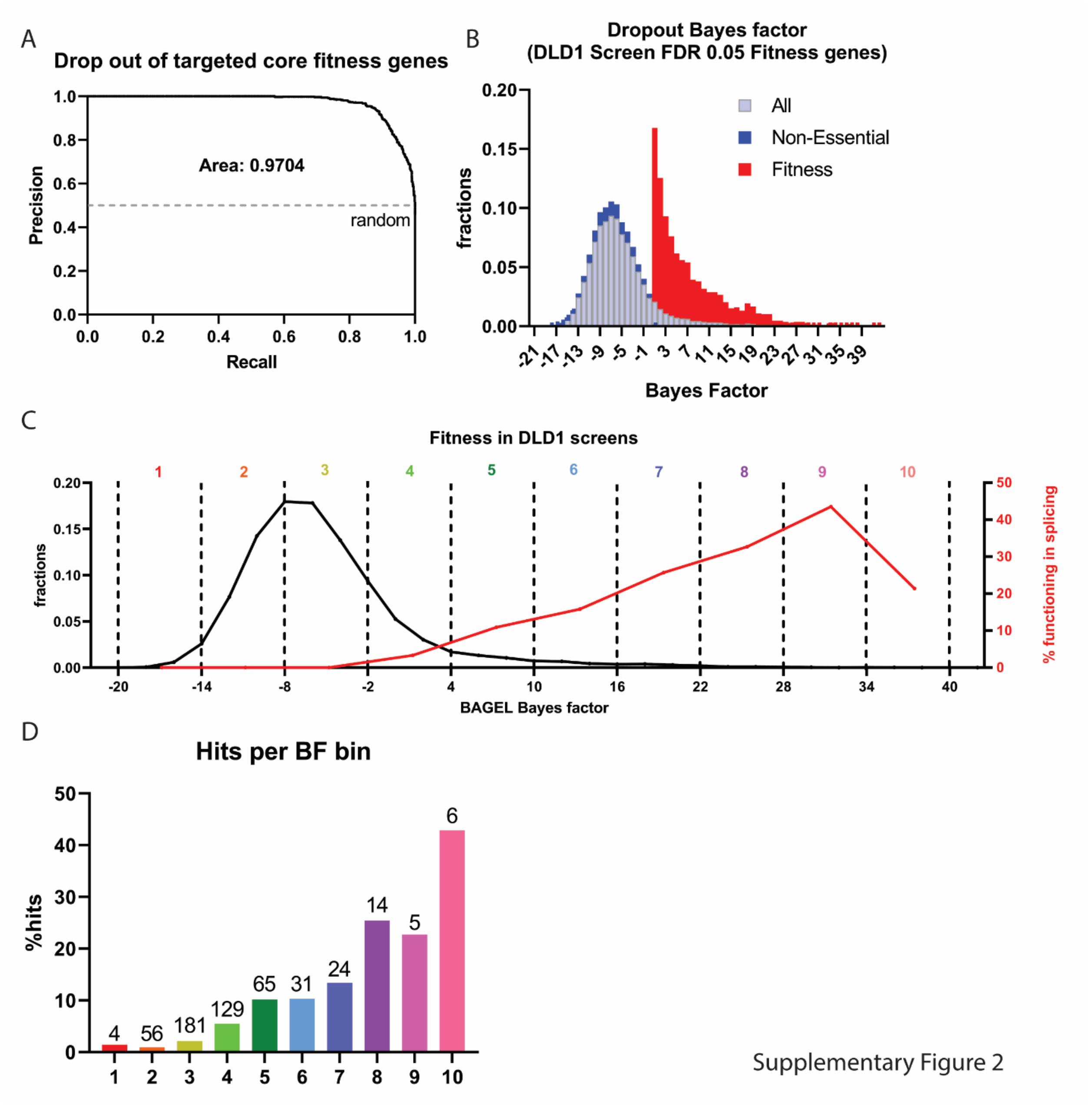


**Supplementary Figure 2: Examination of sgRNA targeting efficiency and effect of fitness on DLD1 CRISPR screens.**

**(A)** Precision-recall plot of core fitness genes (Hart 2017) in DLD1 cells based on BAGEL analysis. The area under the curve score of 0.9704 indicates efficient drop-out of fitness genes indicating that our screens had a good performance considering the shorter screen duration.

**(B)** Bayes factor (BF) distribution of genes targeting core fitness genes (red) or non-essential genes (blue), total library (grey) as determined by BAGEL analysis in unsorted populations of DLD1 Flp-In cells expressing dual fluorescent Numb E9 reporters and transduced with the genome-wide TKOv3 CRISPR library. The BF distributions of core fitness and non-essential genes display little overlap indicating that our CRISPR-Cas9 targeting was successful. Genes with an FDR score <0.05 in the BAGEL analysis were considered fitness.

**(C)** Plot depicting the BF distribution of all genes targeted in the CRISPR screen (black line). Genes were grouped into different BF bins (1-10; as indicated at the top of the plot). The red line indicates the percentage of genes that belong to GO BP related to splicing regulation according to DAVID GO BP analysis (see Fig 8B).

**(D)** Bar chart representing the percentage of CRISPR screen hits in each bin based on BF scores in **(C)** (%hit = number of hits in bin/total number of genes in bin*100). The absolute number of hits in each bin is indicated above each panel.

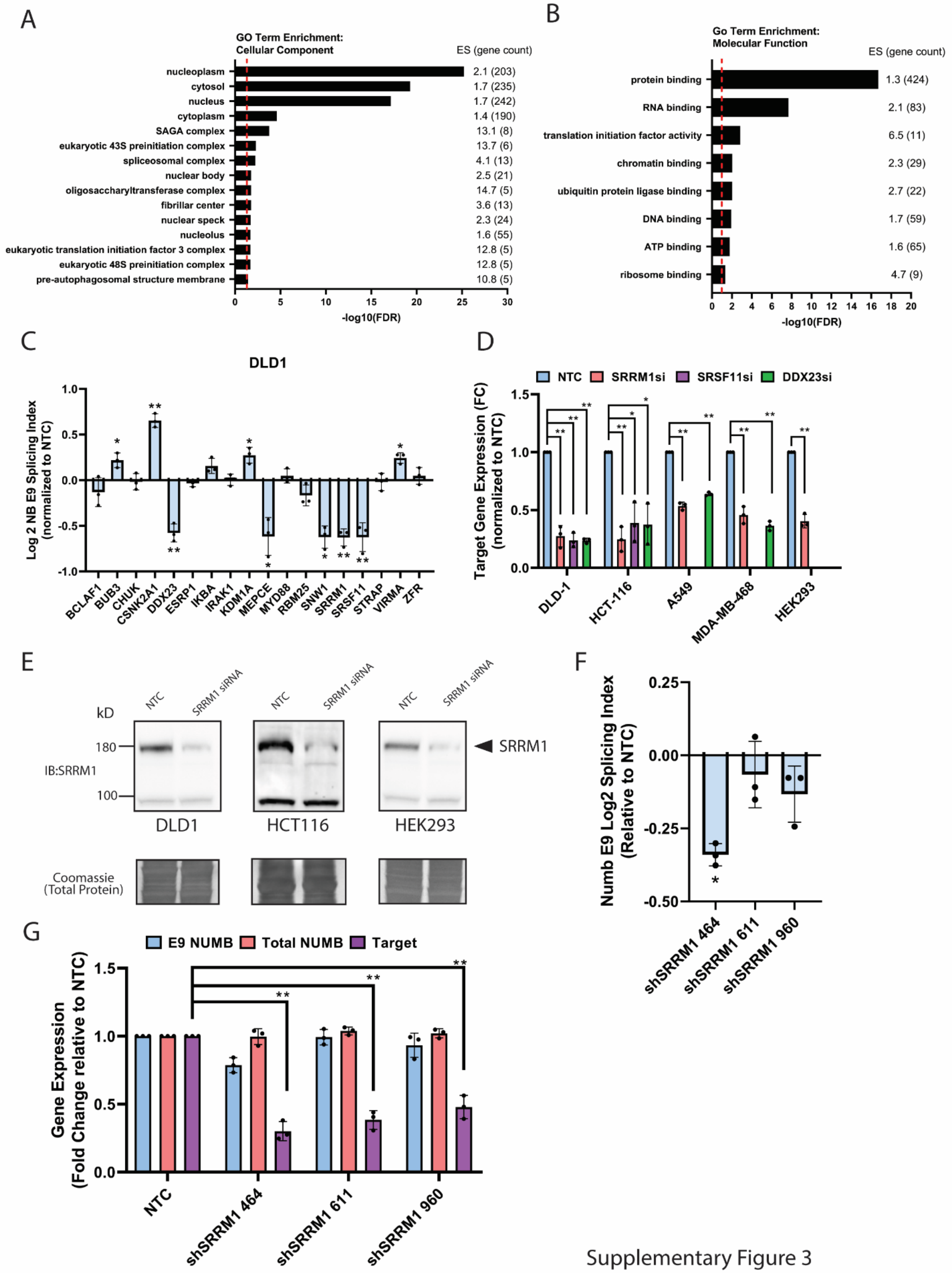


Supplementary Figure 3

**Supplementary Figure 3: Specificity of SRRM1 and SRSF11 knockdown on Numb E9 splicing.**

- **(A)** Cellular component GO term enrichment analysis for the hits from the CRISPR screens. The same hit list as in **(Fig 2A)** was uploaded to the online Database for Annotation, Visualization and Integrated Discovery, DAVID, and clustered using an EASE of 1.0 and medium stringency. The top enriched terms are arranged in order of significance (-log10(FDR) with Enrichment score (ES) values and total hit gene count for each term in between brackets denoted on the right. Only terms with an FDR score < 0.05 are shown. The red dotted line indicates an FDR of 0.1
- **(B)** Similar analysis as in **(A)** for the top molecular function GO term enrichment.
- **(C)** Examination of endogenous Numb E9 splicing index (log2 fold change expression of the Numb+E9 transcript relative to total Numb transcript expression, measured via RT-qPCR.) in DLD1 cells following transfection with siRNAs targeting splicing regulators identified in the CRISPR screen. (n = 3, mean Log2SI ± s.e.m.) All results depicted are relative to non-target control siRNA and significant differences with a p-values < 0.05 indicated with * and p-value < 0.01 with ** according to paired t-test statistical analysis
- **(D)** Measurement of gene expression of the siRNA targeted genes from the validated hits with significant effects on endogenous Numb E9 splicing from **(Fig 2F)** across cell lines relative to non-target control treatment. (n = 3, mean ± s.e.m.)
- **(E)** Immunoblots demonstrate knockdown of SRRM1 relative to non-target control (NTC) following treatment with SRRM1 siRNA in DLD1, HCT116 and HEK293 cells.
- **(F)** Examination of endogenous Numb E9 splicing index in shDLD1 cells targeting SRRM1 (464/611/960) induced with tetracycline for 48hrs. (n = 3, mean Log2SI ± s.e.m.). All results depicted are relative to non-target control shRNA DLD1 cell line with significant differences with a p-values < 0.05 indicated with * according to paired t-test statistical analysis.
- **(G)** Real-time quantitative PCR measuring SRRM1 expression knockdown in response to short hairpin induction relative to shNTC control across 3 biological replicates of shDLD1 cell lines (n = 3, mean ± s.e.m.). (p-value < 0.05 *, p < 0.01 **, p < 0.001***) according to paired t-test statistical analysis

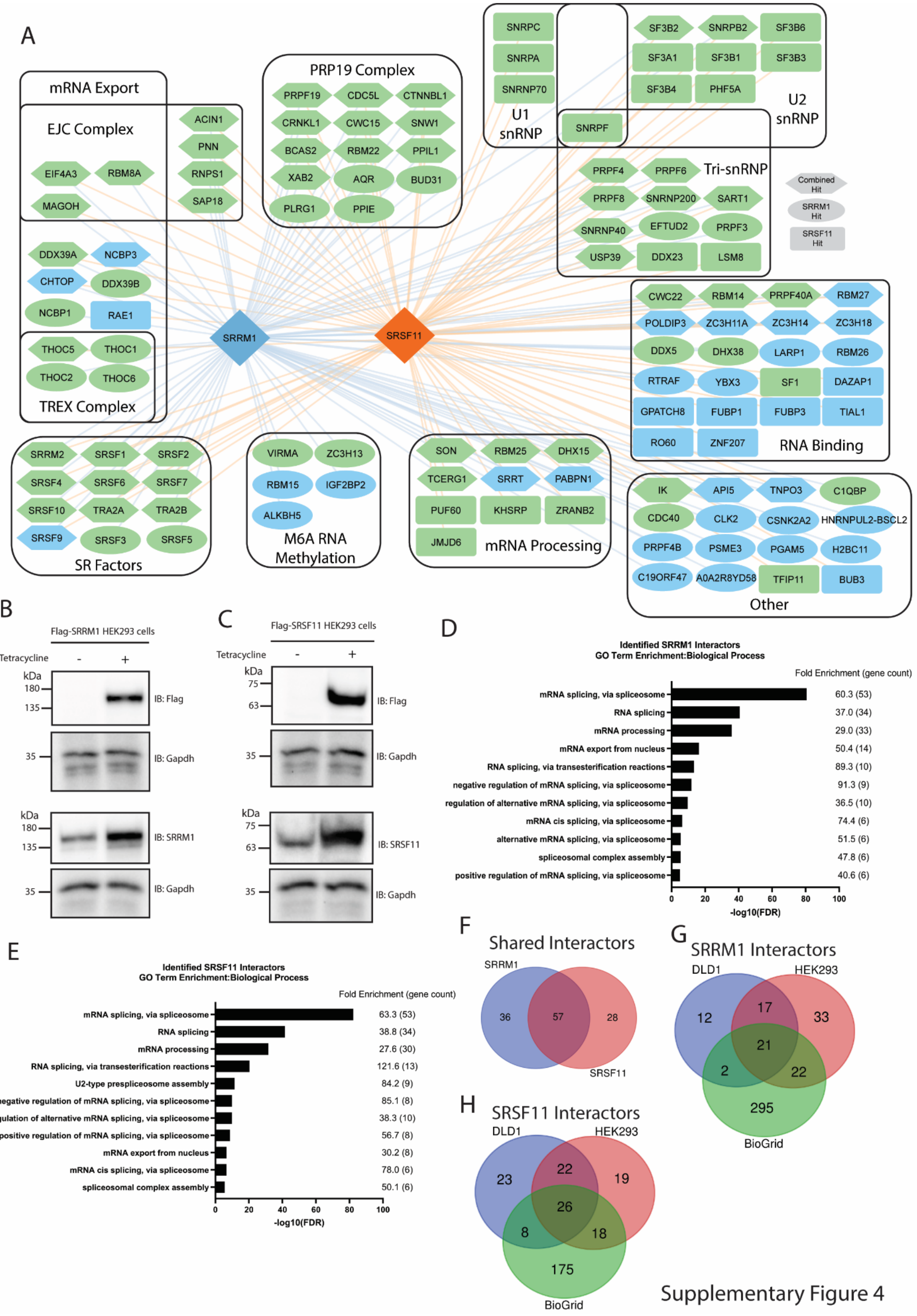


Supplementary Figure 4

**Supplementary Figure 4: Investigating SRRM1/SRSF11 interactome using AP-MS in HEK293 cell line.**

- **(A)** Protein-Protein interactors of SRRM1 (blue edges) and SRSF11 (orange edges) identified via affinity-purification mass spectrometry in HEK293 cells. Interactors with a BFDR ≤ 0.05 are depicted. Unique prey are depicted in ovals for SRRM1 and rectangles for SRSF11 whilst shared prey between the two baits are depicted in hexagons. Green Nodes denote hits that possessed a GOBP term that belongs to RNA splicing and/or mRNA splicing, via spliceosome according to DAVID analysis (https://davidbioinformatics.nih.gov).
- **(B)** Immunoblots demonstrating expression of Flag tagged SRRM1 HEK293 cells ± tetracycline (10ug/ml) induction blotted with either anti-FLAG or anti-SRRM1 antibodies with corresponding anti-Gapdh blots used as a loading control.
- **(C)** Immunoblots demonstrating expression of Flag tagged SRSF11 HEK293 cells ± tetracycline (10ug/ml) induction blotted with anti-FLAG or anti-SRSF11 antibodies with corresponding anti-Gapdh blots used as a loading control.
- **(D)** Biological process GO term enrichment analysis for SRRM1 protein interactors analyzed through DAVID. The top enriched terms are arranged in order of significance (-log10(FDR) with fold enrichment values and total hit gene count for each term in between brackets denoted on the right.
- **(E)** Same as **(B)** for SRSF11 protein interactors.
- **(F)** Venn diagram depicting the number of shared interactors between SRRM1 and SRSF11 identified in HEK293 cells.
- **(G)** Venn diagram depicting the number of shared SRRM1 interactors identified from the DLD1 and HEK293 APMS experiments as well as known SRRM1 interactors obtained from Biogrid database (https://thebiogrid.org/).
- **(H)** Same as **(E)** for SRSF11 interactors.

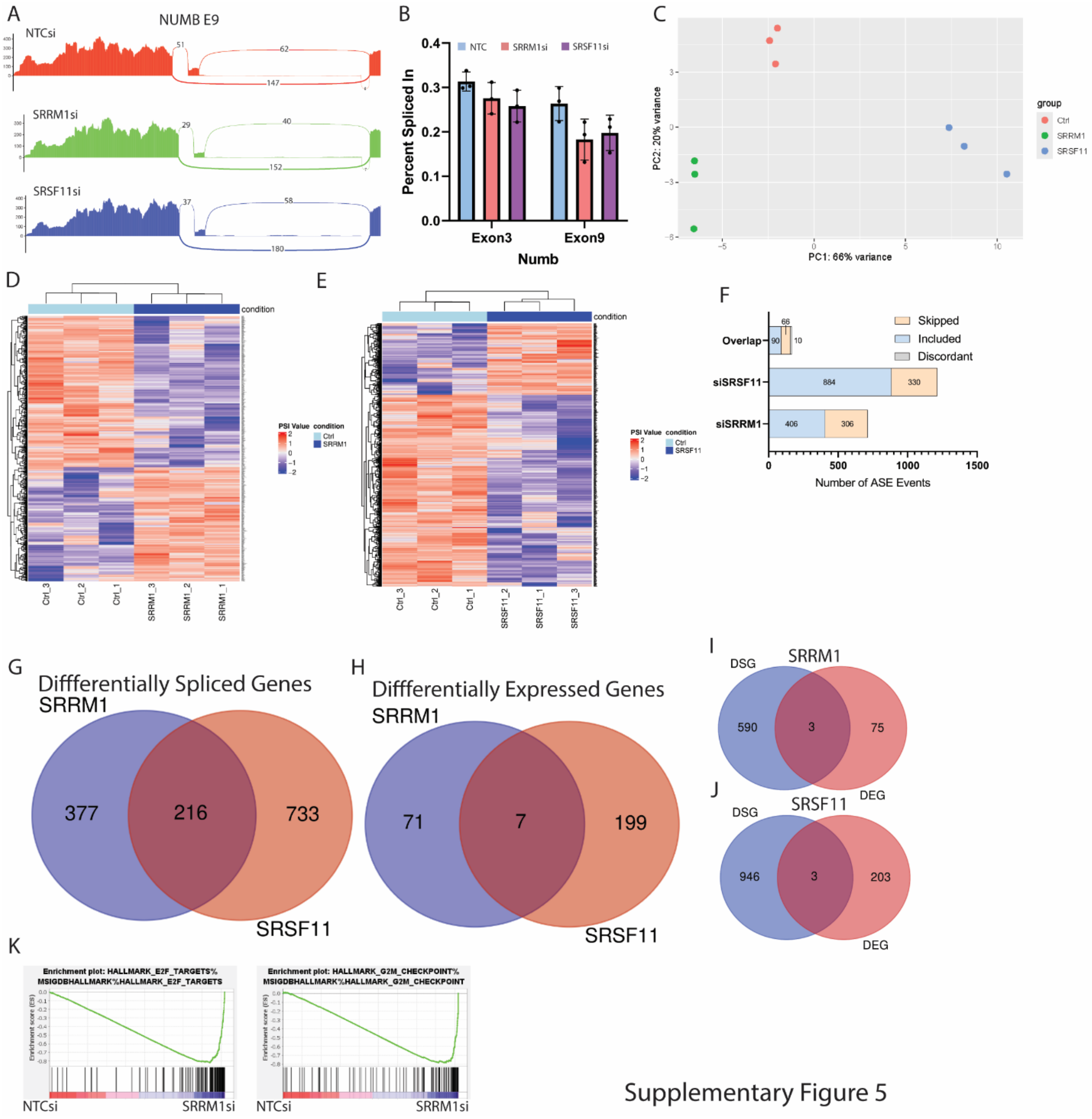


**Supplementary Figure 5: SRRM1 and SRSF11 knockdown results in altered downstream splicing.**

- **(A)** Representative sashimi plots from RNA-seq data depicting Numb E9 inclusion and exclusion reads in siCTRL, siSRRM1 or siSRSF11 conditions.
- **(B)** Related to (**A**) Percent Spliced In (PSI) values of Numb exon 3 and exon 9 in DLD1 cancer cell lines in siCTRL, siSRRM1 or siSRSF11 conditions. N=3 biological replicates
- **(C)** Principal component analyses (PCA) demonstrating clustering of replicates for siCTRL, siSRRM1 and siSRSF11 conditions derived from DLD1 cells.

**(D)** Heatmap of normalized PSI values from SRRM1-regulated alternatively spliced exons (ASEs) in siSRRM1 or siCTRL conditions.

**(E)** Heatmap of normalized PSI values from SRSF11-regulated alternatively spliced exons (ASEs) in siSRSF11 or siCTRL conditions.

**(F)** Bar graph depicting number of SRRM1, SRSF11, or commonly regulated alternatively spliced exons (ASEs) identified in DLD1 cells, representing a summarized version of (**D/E**). Negative PSI shifts in response to silencing are considered Included, depicted in blue, whilst positive PSI shifts are considered skipped depicted in orange.

**(G)** Venn diagram depicting overlap in differentially spliced genes (DSGs) that demonstrated significant dysregulated alternatively spliced exon events (ASE) in the siSRRM1 and siSRSF11 conditions relative to siCTRL, related to Fig 5C.

**(H)** Venn diagram depicting overlap in significantly differentially expressed genes (DEGs) following SRRM1 or SRSF11 silencing in DLD1 colorectal adenocarcinoma cell lines. Data represents 3 biological replicates (siCTRL, siSRRM1 and siSRSF11). FDR <0.05, fold change > 2).

**(I)** Venn diagram depicting overlap between SRRM1-regulated DSGs and DEGs.

**(J)** Venn diagram depicting overlap between SRSF11-regulated DSGs and DEGs

**(K)** Significant terms identified through GSEA of gene expression changes in the siSRRM1 condition.

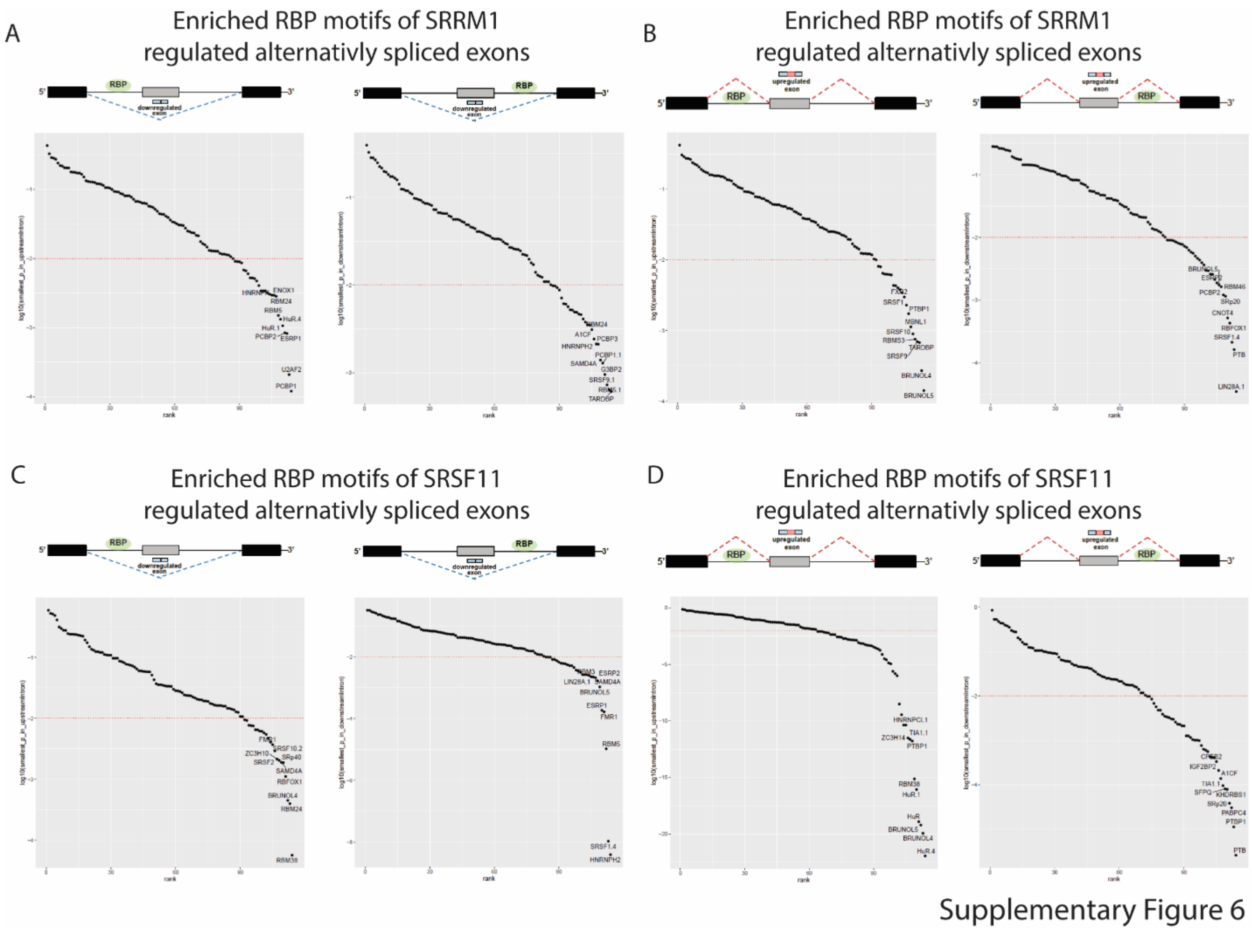


**Supplementary Figure 6: RBP Motif enrichment of dysregulated splicing events downstream of SRRM1 and SRSF11 knockdown identifies potential coordinated regulators.**

**(A-D)** RBP protein motifs are enriched upstream (left) and downstream (right) of SRRM1-regulated (**A**/**B**) or SRSF11-regulated (**C**/**D**) alternatively spliced exons in DLD1 cancer cell line. N=3 biological replicates.

| | Top 20 enriched RBP Motifs identified from SRRM1 regulated ASEs | | | | | | | | | | | |
|---|---|---|---|---|---|---|---|---|---|---|---|---|
| | Upregulated | | | | | | Downregulated | | | | | |
| | Upstream | | | Downstream | | | Upstream | | | Downstream | | |
| | Symbol | Motif | p-value | Symbol | Motif | p-value | Symbol | Motif | p-value | Symbol | Motif | p-value |
| 1 | **CELF5** | TGTGT[GT][GT] | 0.000141966 | **LIN28A** | [CT]GGAGG[AG] | 3.50E-05 | **PCBP1** | C[CT]TTCC | 0.00012066 | **TARDBP** | GAATG[AGT] | 0.00061079 |
| 2 | **CELF4** | [GT]GTGT[GT][GT] | 0.000271713 | **PTBP1** | CTCT[CT][CT] | 0.000163097 | **U2AF2** | TTTTT[CT]C | 0.00020897 | **RBM5** | GA[AG]GG[AT][AG] | 0.00063942 |
| 3 | **TARDBP** | GAATG[AGT] | 0.000678449 | **SRSF1** | GGA[GC]G[AG][ACG] | 0.000213307 | **ESRP1** | TGGTGG | 0.0008239 | **SRSF9** | A[GT]GA[ACG][AC][AG] | 0.00072594 |
| 4 | **SRSF9** | A[GT]GA[ACG][AC][AG] | 0.000696642 | **RBFOX1** | [AT]GCATG[AC] | 0.000431341 | **PCBP2** | CC[CT][CT]CC[ACT] | 0.00084114 | **G3BP2** | AGGAT[AGT][AG] | 0.00095485 |
| 5 | **RBMS3** | [ACT]ATATA | 0.000753211 | **CNOT4** | GACAGA | 0.000520308 | **ELAVL1** | TT[GT][AG]TTT | 0.00106302 | **PCBP1** | CC[AT][AT][ACT]CC | 0.0012856 |
| 6 | **SRSF10** | A[AG]AG[AG][AG][AG] | 0.000897858 | **SRSF3** | [AT]C[AT][AT]C | 0.001154567 | **ELAVL1** | TT[AGT]TTTT | 0.00132226 | **SAMD4A** | GC[GT]GG[ACT][AC] | 0.00139476 |
| 7 | **MBNL1** | GCTTGC | 0.001131192 | **PCBP2** | CC[CT][CT]CC[ACT] | 0.001213541 | **RBM5** | [GC]AAGG[AG]G | 0.00149919 | **PCBP3** | TTT[CT]CC | 0.00211535 |
| 8 | **PTBP1** | [ACT][CT]TTT[CT]T | 0.001736408 | **RBM46** | [AG]AT[GC]A[AT][AGT] | 0.001634591 | **RBM24** | [AT]G[AT]GTG[AGT] | 0.00283962 | **HNRNPH2** | GGGAGGG | 0.00212607 |
| 9 | **SRSF1** | GG[AG]GGA[ACG] | 0.002296508 | **CELF5** | TGTGT[GT][GT] | 0.001753549 | **ENOX1** | [ACT][AG][GT]ACAG | 0.00294347 | **A1CF** | [AT]TAATT[AG] | 0.00242162 |
| 10 | **FXR2** | [AGT]GAC[AG][AG][AG] | 0.002989684 | **ESRP2** | TGGG[AG]A[AGT] | 0.001887738 | **HNRNPK** | CCA[AT][AC]CC | 0.00294998 | **RBM24** | [AT]G[AT]GTG[AGT] | 0.00310676 |
| 11 | **SRSF2** | GGAG[AT][AGT] | 0.003451027 | **SRSF9** | A[GT]GA[ACG][AC][AG] | 0.002133406 | **RBFOX1** | [AT]GCATG[AC] | 0.00304462 | **PTBP1** | CTCT[CT][CT] | 0.00349484 |
| 12 | **RBM5** | [GC]AAGG[AG]G | 0.003811326 | **SRSF5** | [CT][AG]C[AG][GT][AC] | 0.002589512 | **PTBP1** | [ACT][CT]TTT[CT]T | 0.00317937 | **ESRP1** | TGGTGG | 0.00353876 |
| 13 | **IGF2BP2** | [ACG][AC]A[ACT][AT]CA | 0.004032612 | **SRSF3** | CTC[GT]TC[CT] | 0.002602977 | **KHDRBS2** | [AG]ATAAA[AC] | 0.00340442 | **RBM6** | [ACT]ATCCA[AG] | 0.00378077 |
| 14 | **HNRNPK** | CCA[AT][AC]CC | 0.004330962 | **ELAVL1** | TT[GT][AG]TTT | 0.002949864 | **SART3** | A[AG]AAAA[AC] | 0.00340442 | **U2AF2** | TTTTT[CT]C | 0.00409933 |
| 15 | **SRSF6** | [CT][CT][AT]C[AT][GC]G | 0.004360756 | **RBM5** | [GC]AAGG[AG]G | 0.002977725 | **PABPC1** | A[AG]AAAA[AC] | 0.00340442 | **RBMS1** | [GT]ATATA[GC] | 0.00459963 |
| 16 | **PCBP1** | CC[AT][AT][ACT]CC | 0.006142482 | **CELF4** | [GT]GTGT[GT][GT] | 0.00363661 | **ZC3H14** | TTT[AGT]TTT | 0.00406555 | **FXR1** | A[CT]GAC[AG] | 0.00466263 |
| 17 | **SNRPA** | [AT]TGCAC[AG] | 0.006215263 | **SRSF1** | GGAGGA | 0.003946678 | **DAZAP1** | TAG[GT][AT][AT][AG] | 0.00472559 | **PABPN1** | A[AG]AAGA | 0.00490175 |
| 18 | **SRSF3** | [AT]C[AT][AT]C | 0.006255488 | **RBM24** | [AT]G[AT]GTG[AGT] | 0.004358436 | **SRSF1** | G[AG]AGGA | 0.00511327 | **SRSF9** | [GT]G[AG][AT]G[GC][AC] | 0.00493256 |
| 19 | **MSI1** | TAGT[AT][AG]G | 0.00629408 | **SRSF1** | GG[AG]GGA[ACG] | 0.004646407 | **SRSF1** | GG[AG]GGA[ACG] | 0.00523248 | **KHDRBS1** | ATAAAA[ACG] | 0.00517529 |
| 20 | **TRA2B** | GAAAGAA | 0.006484067 | **QKI** | ACTAAC[ACG] | 0.005042378 | **TIA1** | TTTTT[CGT][GT] | 0.00525077 | **SRSF5** | [CT][AG]C[AG][GT][AC] | 0.00522861 |

**Supplementary Table1: Top 20 Enriched RBP motifs identified from SRRM1-regulated ASEs**

| | Top 20 enriched RBP Motifs identified from SRSF11 regulated ASEs | | | | | | | | | | | |
|---|---|---|---|---|---|---|---|---|---|---|---|---|
| | Upregulated | | | | | | Downregulated | | | | | |
| | Upstream | | | Downstream | | | Upstream | | | Downstream | | |
| | Symbol | Motif | p-value | Symbol | Motif | p-value | Symbol | Motif | p-value | Symbol | Motif | p-value |
| 1 | **ELAVL1** | TT[AT]GTTT | 1.13E-22 | PTBP1 | CTCT[CT][CT] | 1.12E-05 | RBM38 | [GT][GT]GTGT[GT] | 5.67E-05 | HNRNPH2 | GGGAGGG | 3.92E-09 |
| 2 | **CELF4** | [GT]GTGT[GT][GT] | 1.16E-20 | PTBP1 | [ACT][CT]TTT[CT]T | 1.65E-12 | RBM24 | [AT]G[AT]GTG[AGT] | 0.000399 | SRSF1 | GGA[GC]G[AG][ACG] | 1.07E-08 |
| 3 | **CELF5** | TGTGT[GT][GT] | 6.12E-20 | PABPC4 | AAAAAA[AG] | 0.015522487 | CELF4 | [GT]GTGT[GT][GT] | 0.000449 | RBM5 | GA[AG]GG[AT][AG] | 1.05E-05 |
| 4 | **ELAVL1** | TT[GT][AG]TTT | 1.22E-19 | SRSF3 | [AT]C[AT][AT]C | 0.002770535 | RBFOX1 | [AT]GCATG[AC] | 0.001108 | FMR1 | [GT]GACA[AG]G | 0.000165 |
| 5 | **ELAVL1** | TTT[AG][GT]TT | 8.55E-17 | KHDRBS1 | TAAAA[ACG][ACG] | 0.000239442 | SRSF5 | [CT][AG]C[AG][GT][AC] | 0.001879 | ESRP1 | TGGTGG | 0.000185 |
| 6 | **RBM38** | [GT][GT]GTGT[GT] | 7.42E-16 | SFPQ | [GT]T[AG][AG]T[GT][GT] | 3.19E-05 | SAMD4A | GC[GT]GG[ACT][AC] | 0.001899 | CELF5 | TGTGT[GT][GT] | 0.001058 |
| 7 | **PTBP1** | [ACT][CT]TTT[CT]T | 1.65E-12 | TIA1 | TTTTT[CGT][GT] | 2.35E-12 | SRSF2 | GGAG[AT][AGT] | 0.002048 | SAMD4A | GC[GT]GG[ACT][AC] | 0.001664 |
| 8 | **TIA1** | TTTTT[CGT][GT] | 2.35E-12 | A1CF | [AT]TAATT[AG] | 0.006197623 | ZC3H10 | [GC][GC]AGCG[AC] | 0.002143 | ESRP2 | TGGG[AG]A[AGT] | 0.002105 |
| 9 | **ZC3H14** | TTT[AGT]TTT | 3.11E-12 | IGF2BP2 | [ACG][AC]A[ACT][AT]CA | 0.00044146 | SRSF10 | AGAGA[ACG][ACG] | 0.00293 | RBM3 | [AG]A[AGT]AC[GT]A | 0.002205 |
| 10 | **HNRNPC** | [ACT]TTTTT[GT] | 4.32E-11 | CPEB2 | C[ACT]TTTTT | 0.000289999 | FMR1 | [GT]GACA[AG]G | 0.003696 | LIN28A | [ACT]GGAG[AT]A | 0.002266 |
| 11 | **HNRNPCL1** | [ACT]TTTTT[GT] | 4.32E-11 | HNRNPL | A[AC]A[CT]A[AC]A | 0.002728976 | SRSF1 | GG[AG]GGA[ACG] | 0.004056 | SRSF9 | [GT]G[AG][AT]G[GC][AC] | 0.002504 |
| 12 | **ELAVL1** | TT[AGT]TTTT | 3.55E-10 | ELAVL1 | TT[GT][AG]TTT | 1.22E-19 | RBMS1 | [GT]ATATA[GC] | 0.004717 | HNRNPA1L2 | [AGT]TAGGG[AT] | 0.002608 |
| 13 | **ZNF638** | [CGT]GTT[GC][GT]T | 3.14E-09 | PABPC3 | [AG]AAAAC[AC] | 0.013396639 | SRSF10 | AGAGA[AG][AG] | 0.005459 | HNRNPA1 | [AGT]TAGGG[AT] | 0.002608 |
| 14 | **CPEB4** | TTTTTT | 1.00E-06 | SRSF3 | CTC[GT]TC[CT] | 0.033120803 | PCBP2 | CC[CT][CT]CC[ACT] | 0.005772 | HNRNPA2B1 | [AGT]TAGGG[AT] | 0.002608 |
| 15 | **TIA1** | [AT]TTTTT[CGT] | 1.54E-06 | RBM46 | [AG]AT[GC]A[AT][AGT] | 0.004293434 | SRSF1 | G[AG]AGGA | 0.006117 | SRSF1 | AGGA[GC][AC] | 0.003167 |
| 16 | **RBMS3** | [ACT]ATATA | 2.45E-06 | RBM41 | [AT]TAC[AT]T[GT] | 0.001412207 | SRSF1 | GGAGGA | 0.006185 | RBM8A | [AG][CT]GCGC[CGT] | 0.003461 |
| 17 | **PTBP1** | CTCT[CT][CT] | 1.12E-05 | TIA1 | [AT]TTTTT[CGT] | 1.54E-06 | YBX1 | AACATC[AGT] | 0.006476 | CELF4 | [GT]GTGT[GT][GT] | 0.003718 |
| 18 | **RALY** | TTTTTT[CGT] | 1.37E-05 | KHDRBS3 | ATAAA[ACG] | 0.000710999 | SRSF9 | [GT]G[AG][AT]G[GC][AC] | 0.0065 | RBM4 | GCGCG[GC][GC] | 0.004519 |
| 19 | **ELAVL1** | TTTTTT[GT] | 2.54E-05 | HNRNPC | [ACT]TTTTT[GT] | 4.32E-11 | FUS | CGCGC | 0.007508 | YBX2 | AACA[AT]C[AGT] | 0.005138 |
| 20 | **SFPQ** | [GT]T[AG][AG]T[GT][GT] | 3.19E-05 | HNRNPCL | [ACT]TTTTT[GT] | 4.32E-11 | RBMS3 | [AC]TATA[GT][AC] | 0.007609 | PCBP1 | C[CT]TTCC | 0.005181 |

**Supplementary Table 2: Top 20 Enriched RBP motifs identified from SRSF11-regulated ASEs**

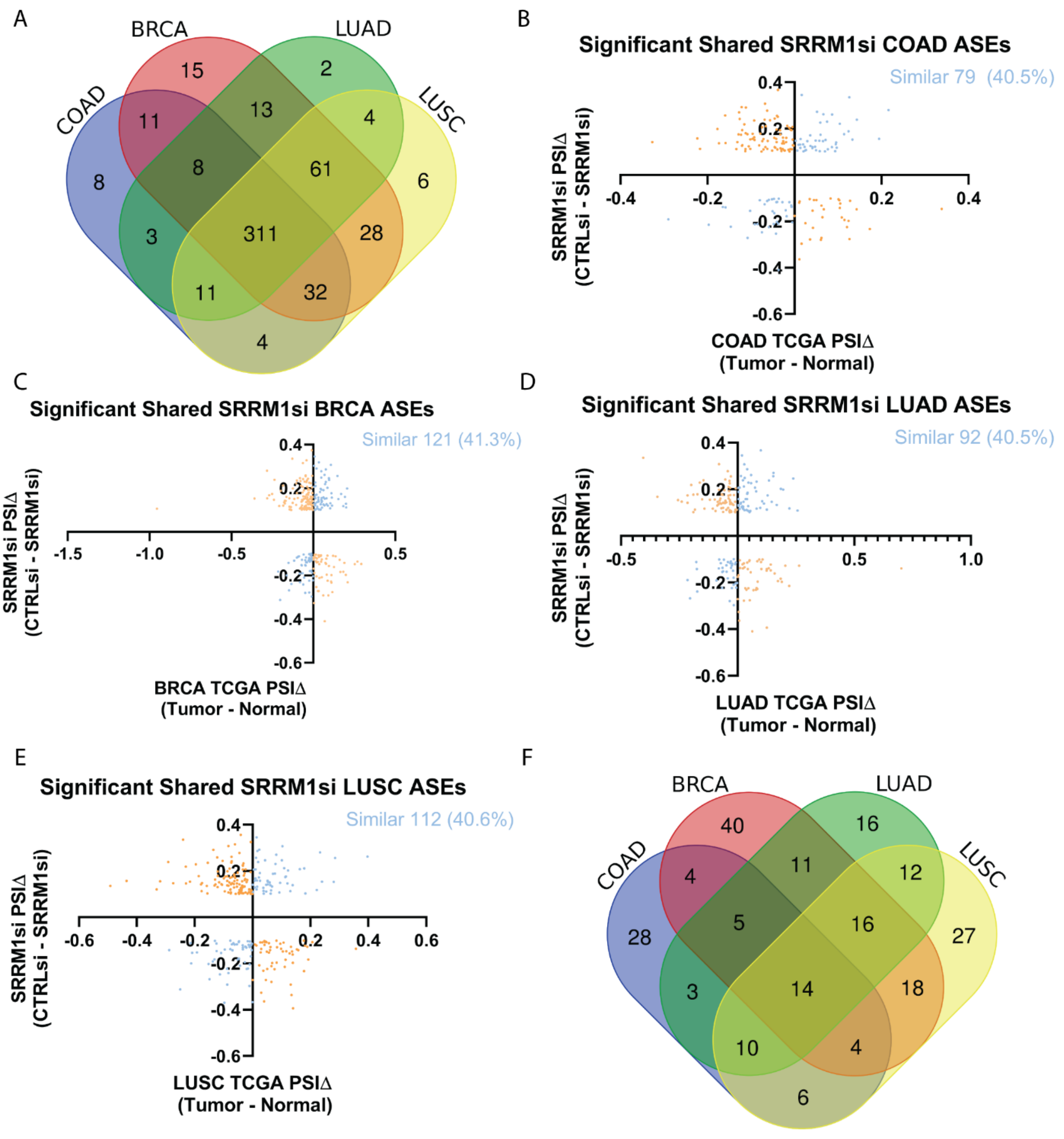


Supplementary Figure 7

**Supplementary Figure 7: Comparisons of SRRM1si ASEs to significant TCGA PSI shifts**

- **(A)** Common dysregulated spliced genes (DSGs) that exhibit significant PSI shifts from TCGA patient samples (BRCA, LUAD, LUSC, COAD) and identified SRRM1-regulated DSGs
- **(B)** Comparison of SRRM1-regulated alternatively spliced events (ASEs) PSIΔ (CTRLsi – SRRM1si) to Mean PSIΔ (Tumor – Normal) from TCGA COAD patient samples.

**(C)** Comparison of SRRM1-regulated ASEs PSIΔ (CTRLsi – SRRM1si) to Mean PSIΔ (Tumor – Normal) from TCGA BRCA patient samples.

**(D)** Comparison of SRRM1-regulated ASEs PSIΔ (CTRLsi – SRRM1si) to Mean PSIΔ (Tumor – Normal) from TCGA LUAD patient samples.

**(E)** Comparison of SRRM1-regulated ASEs PSIΔ (CTRLsi – SRRM1si) to Mean PSIΔ (Tumor – Normal) from TCGA LUSC patient samples.

**(F)** Exon matched splicing events that demonstrated similar shifts in PSI in response to SRRM1 knockdown when compared to changes in PSI between tumor and normal TCGA samples from COAD, BRCA, LUAD and LUSC patients.

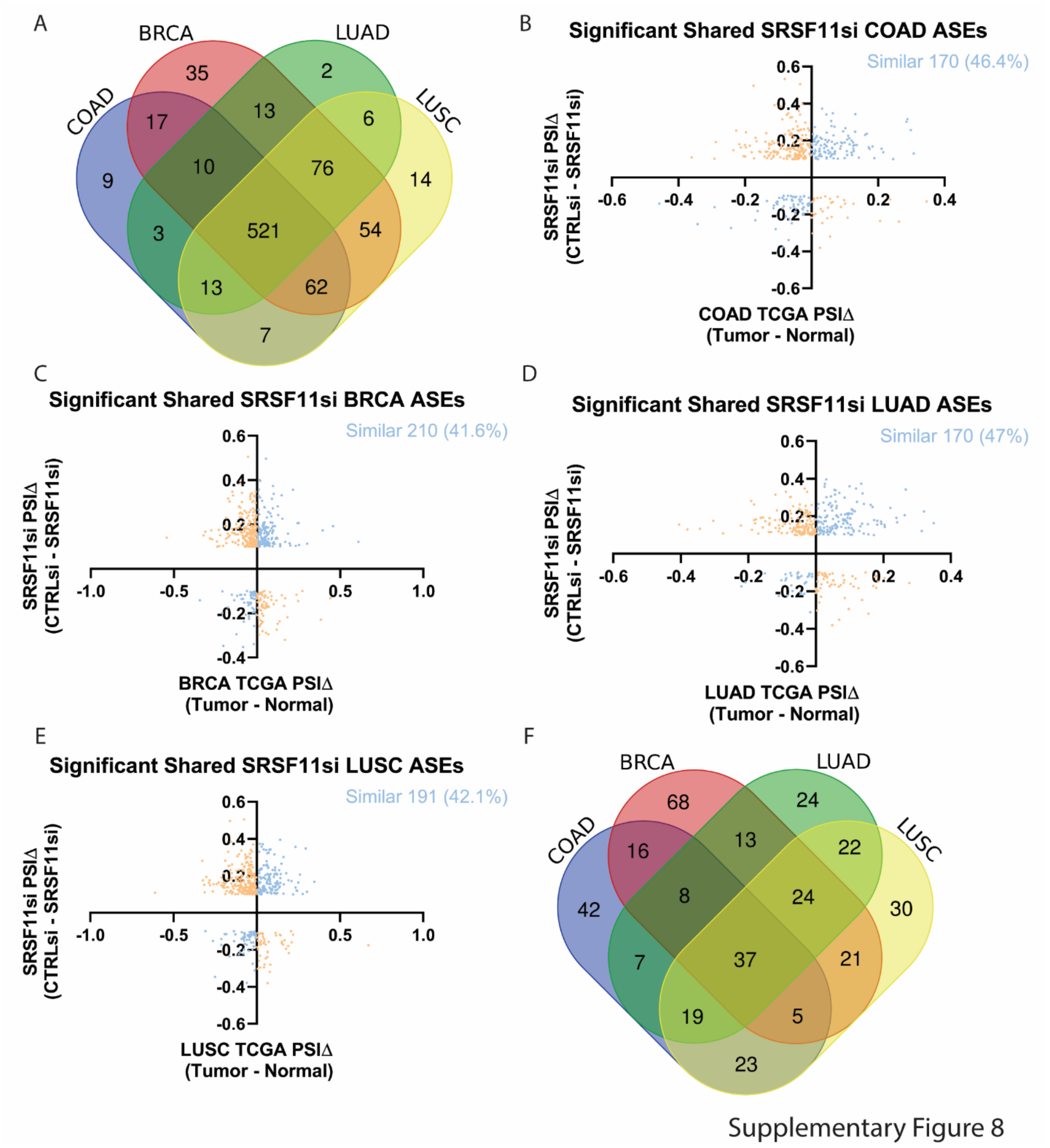


**Supplementary Figure 8: Comparisons of SRSF11si ASEs to significant TCGA PSI shifts**

**(A)** Common DSGs that exhibit significant PSI shifts from TCGA patient samples (BRCA, LUAD, LUSC, COAD) and identified SRSF11-regulated DSGs

**(B)** Comparison of SRSF11-regulated ASEs PSIΔ (CTRLsi – SRSF11si) to Mean PSIΔ (Tumor – Normal) from TCGA COAD patient samples.

**(C)** Comparison of SRSF11-regulated ASEs PSIΔ (CTRLsi – SRSF11si) to Mean PSIΔ (Tumor – Normal) from TCGA BRCA patient samples.

**(D)** Comparison of SRSF11-regulated ASEs PSIΔ (CTRLsi – SRSF11si) to Mean PSIΔ (Tumor – Normal) from TCGA LUAD patient samples.

**(E)** Comparison of SRSF11-regulated ASEs PSIΔ (CTRLsi – SRSF11si) to Mean PSIΔ (Tumor – Normal) from TCGA LUSC patient samples.

**(F)** Exon matched splicing events that demonstrated similar shifts in PSI in response to SRSF11 knockdown when compared to changes in PSI between tumor and normal TCGA samples from COAD, BRCA, LUAD and LUSC patients.

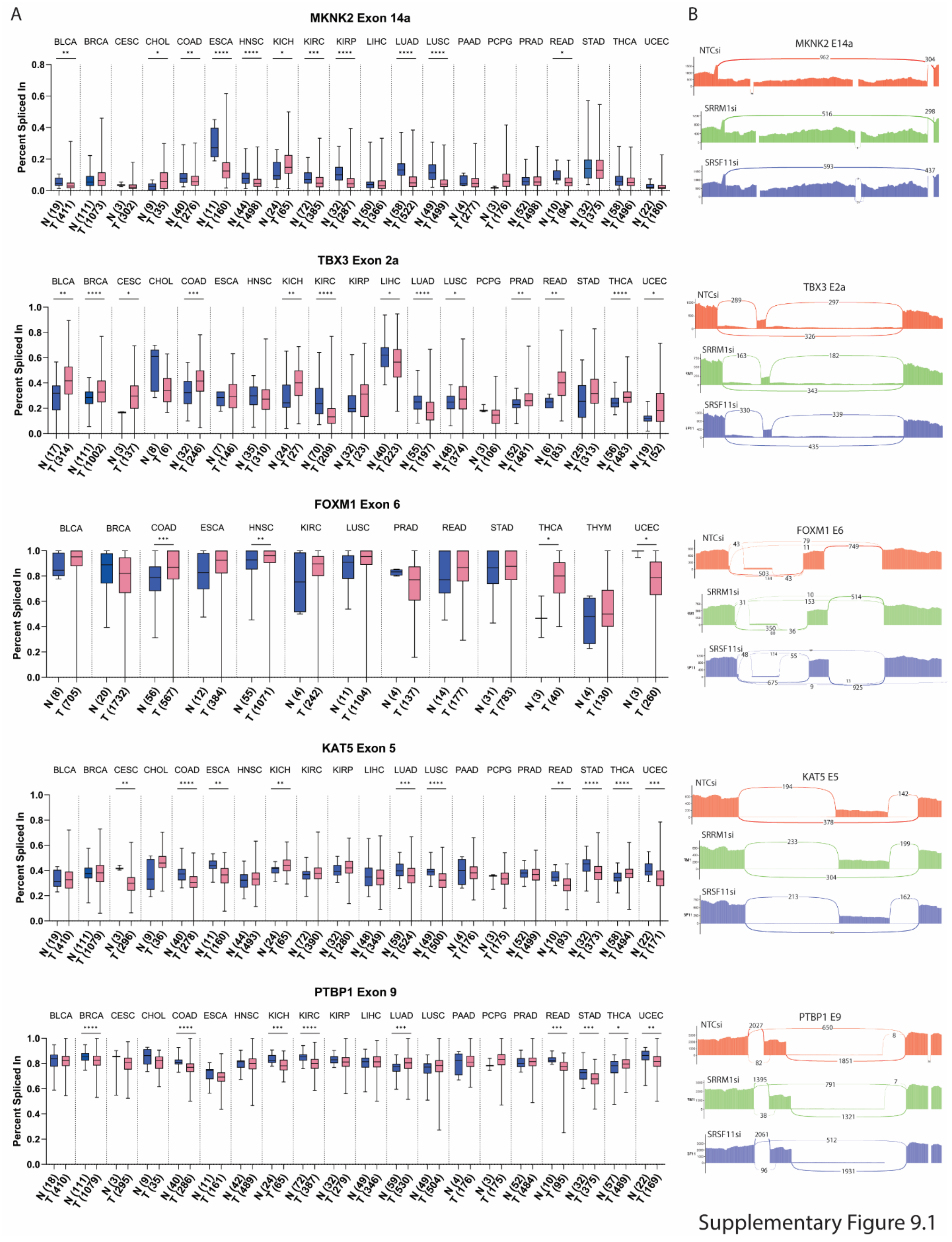


Supplementary Figure 9.1

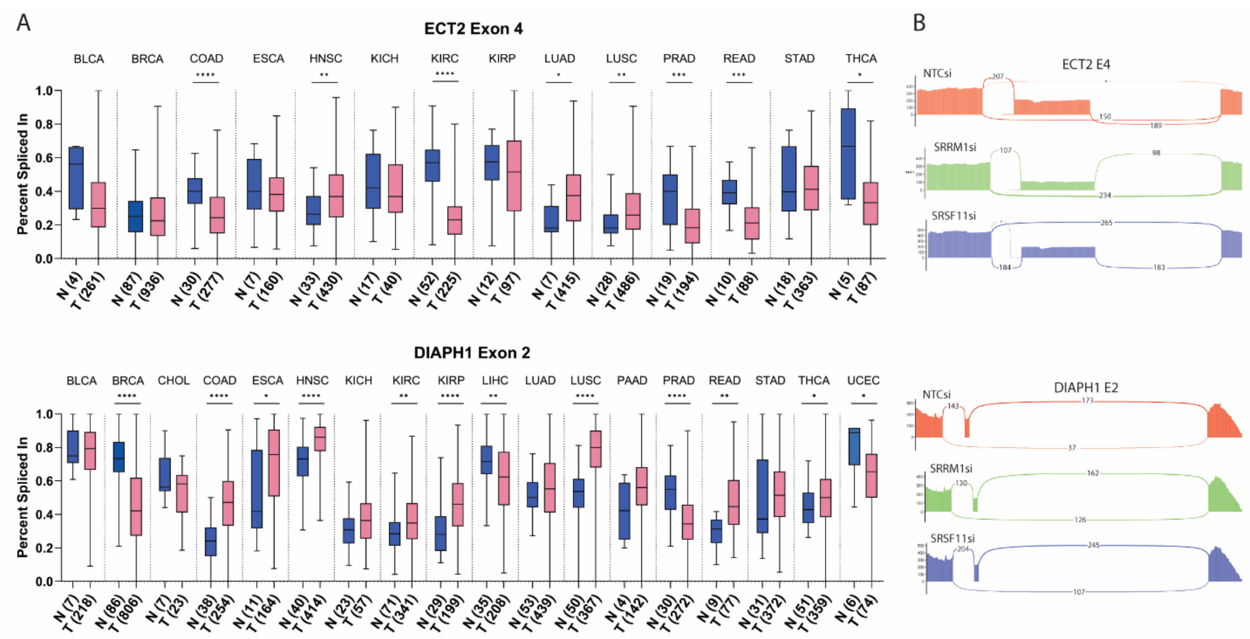


Supplementary Figure 9.2

**Supplementary Figure 9: Examination of MKNK2, FOXM1, TB3, KAT5, PTBP1, ECT2, and DIAPH1 ASEs across TCGA samples and DLD1 RNA-seq data**

**(A)** MKNK2 E14, FOXM1E6, TBX E2a, KAT5 E5, PTBP1 E9, ECT2 E4, and DIAPH1 E2 PSI values in tumor and patient matched normal samples across TCGA dataset obtained from (Kahles et al. 2018) (T=Tumor, N= Normal). A Wilcoxon test was performed comparing tumor and the normal samples. *$p \leq 0.05$; **$p \leq 0.01$;***$p \leq 0.001$; ****$p \leq 0.0001$; the number of samples is in brackets .

**(B)** Sashimi plots of splicing events examined in **(A)** in NTCsi, SRRM1si and SRSF11si DLD1 RNA-seq data.

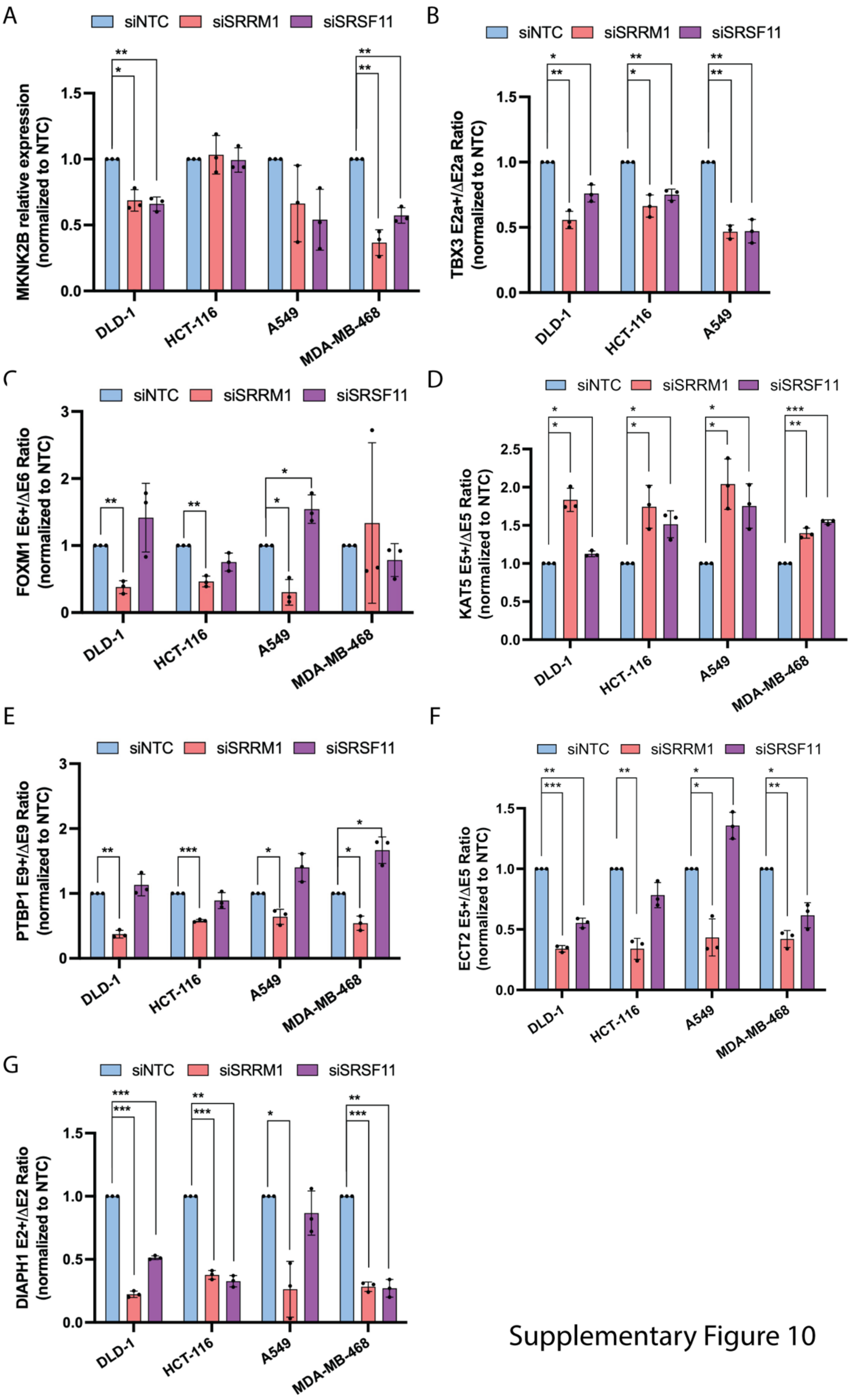


Supplementary Figure 10

**Supplementary Figure 10: Validation of dysregulated splicing events across cancer cell lines**

Validation of SRRM1 and SRSF11 knockdown effect on (**A**) relative MKNK2B expression or shifts in isoform expression ratios of (**B**) TBX3 exon 2a, (**C**) FOXM1 exon 6, (**D**) KAT5 exon 5, (**E**) PTBP1 exon 9, (**F**) ECT2 exon 4, and (**G**) DIAPH1 exon 2 in DLD1, A549, MDA-MB-468, and HCT116 cancer cell lines. ($n = 3$, mean ± SD). (p-value $< 0.05$ *, $p < 0.01$ **, $p < 0.001$***) according to paired t-test statistical analysis.

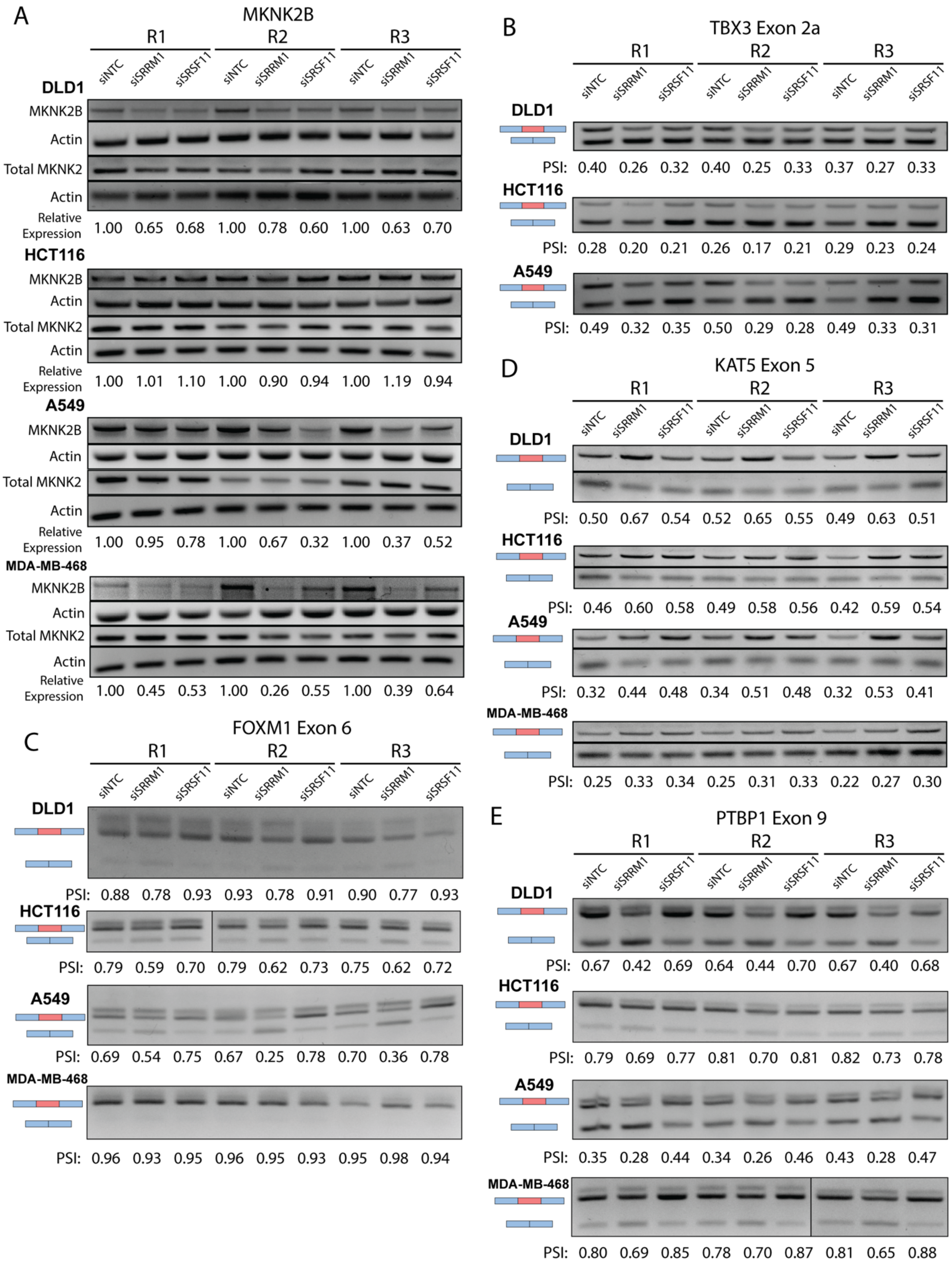


Supplementary Figure 11.1

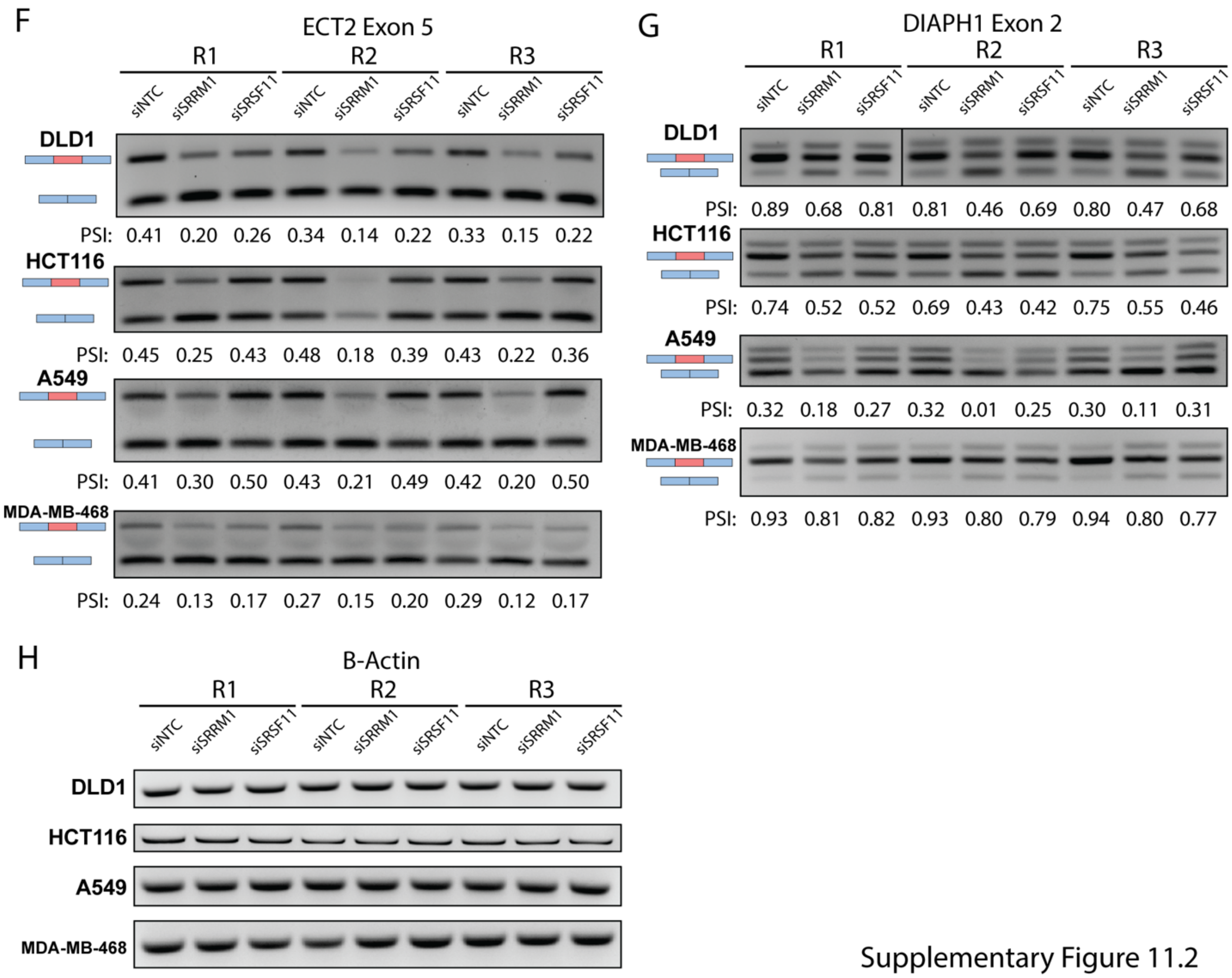


Supplementary Figure 11.2

**Supplementary Figure 11: Validation of dysregulated splicing events across cancer cell lines**

Validation of SRRM1 and SRSF11 knockdown effect on (**A**) relative MKNK2B expression or shifts in isoform expression ratios of (**B**) TBX3 exon 2a, (**C**) FOXM1 exon 6, (**D**) KAT5 exon 5, (**E**) PTBP1 exon 9 (**F**) ECT2 exon 4 (**G**) DIAPH1 exon 2 in DLD1, A549, MDA-MB-468, and HCT116 cancer cell lines. Relative MKNK2B expression was quantified relative to actin loading control (**H**) and then normalized to NTC. Percent spliced in (PSI) was quantified as total inclusion band intensity relative to inclusion and exclusion band intensity.

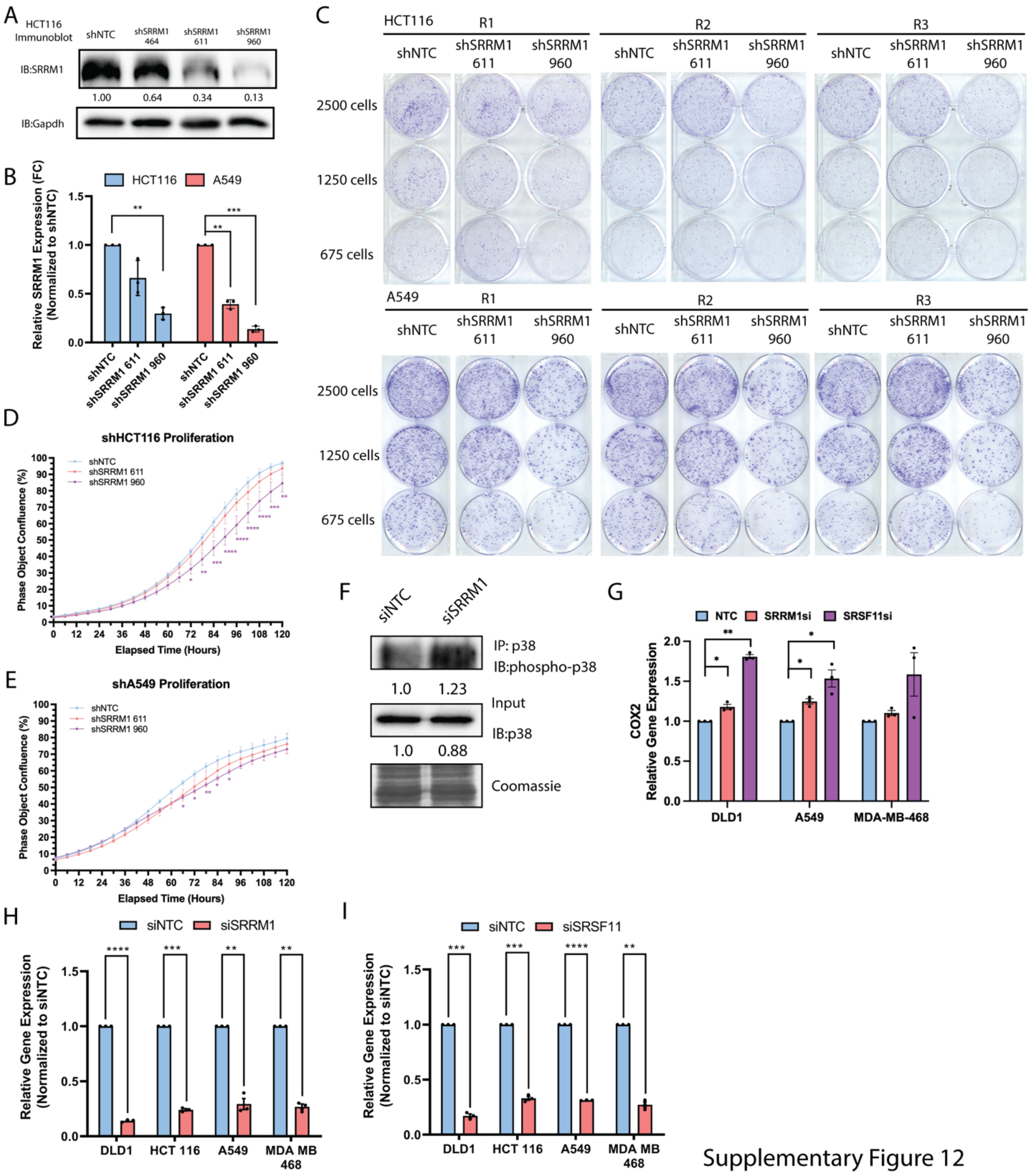


**Supplementary Figure 12: Effect of SRRM1 knockdown on HCT116 colony formation, migration and invasion.**

**(A)** Immunoblots demonstrated relative SRRM1 expression in HCT116 cell lines expressing stet-inducible short hairpin targeting SRRM1 (464/611/960) or a non-target control (NTC) following induction with tetracycline for 48hrs. Numbers under blots denote quantification of relative band intensity normalized to NTC control.

**(B)** Real-time quantitative PCR measuring SRRM1 expression knockdown in response to short hairpin induction relative to shNTC control across 3 biological replicates of shHCT116 and shA549 cell lines ($n = 3$, mean ± s.e.m.). (p-value < 0.05 *, p < 0.01 **, p < 0.001***) according to paired t-test statistical analysis.

**(C)** Representative images from colony formation assay conducted on tetracycline-inducible HCT116 and A549 SRRM1 short hairpin cell lines (shSRRM1 611/960) relative to a non-target control line (NTC) seeded at either 2500, 1250 and 675 cells per well and cultured for 14 days (HCT116) or 10 days (A549) under tetracycline induction before crystal violet staining and imaging

**(D)** Quantification of proliferation assays conducted on HCT116 short hairpin cell lines measuring phase area confluence using an Incucyte live cell imaging system. Cells were cultured at low confluency under tetracycline induction and allowed to grow for 120 hours.

**(E)** Same as **(C)** conducted on A549 short hairpin cell lines.

**(F)** Immunoblots demonstrating phospho-p38 (Thr180/Tyr182) levels in DLD1 cell lines treated with siRNA targeting SRRM1 compared to non-target control following immunoprecipitation of p38 as well as total p38 expression levels. Numbers under blots denote quantification of relative band intensity utilizing Bio-Rad imaging software normalized to NTC control

**(G)** Real-time quantitative PCR measuring COX2 expression in DLD1, A549, and MDA-MB-468 cells treated with SRRM1 or SRSF11 siRNA relative to non-target control (NTC) treatment across 3 biological replicates. Relative fold change expression values were normalized to NTC control. Significant differences with a p-values < 0.05 = *, p-values < 0.01 = **.

**(H)** Confirmation of SRRM1 target knockdown in response to siSRRM1 treatment relative to siNTC control across 3 biological replicates. ($n = 3$, mean ± s.e.m.). (p-value < 0.05 *, p < 0.01 **, p < 0.001***) according to paired t-test statistical analysis.

**(I)** Confirmation of SRSF11 target knockdown in response to siSRSF11 treatment relative to siNTC control across 3 biological replicates. ($n = 3$, mean ± s.e.m.). (p-value < 0.05 *, p < 0.01 **, p < 0.001***) according to paired t-test statistical analysis.
